# Regulation of adenylyl cyclase 5 in striatal neurons confers the ability to detect coincident neuromodulatory signals

**DOI:** 10.1101/597096

**Authors:** Neil J. Bruce, Daniele Narzi, Daniel Trpevski, Siri Camee van Keulen, Anu G. Nair, Ursula Röthlisberger, Rebecca C. Wade, Paolo Carloni, Jeanette Hellgren Kotaleski

## Abstract

Long-term potentiation and depression of synaptic activity in response to stimuli is a key factor in reinforcement learning. Strengthening of the corticostriatal synapses depends on the second messenger cAMP, whose synthesis is catalysed by the enzyme adenylyl cyclase 5 (AC5), which is itself regulated by the stimulatory Gα_olf_ and inhibitory Gα_i_ proteins. AC isoforms have been suggested to act as coincidence detectors, promoting cellular responses only when convergent regulatory signals occur close in time. However, the mechanism for this is currently unclear, and seems to lie in their diverse regulation patterns. Despite attempts to isolate the ternary complex, it is not known if Gα_olf_ and Gα_i_ can bind to AC5 simultaneously, nor what activity the complex would have. Using protein structure-based molecular dynamics simulations, we show that this complex is stable and inactive. These simulations, along with Brownian dynamics simulations to estimate protein association rates constants, constrain a kinetic model that shows that the presence of this ternary inactive complex is crucial for AC5’s ability to detect coincident signals, producing a synergistic increase in cAMP. These results reveal some of the prerequisites for corticostriatal synaptic plasticity, and explain recent experimental data on cAMP concentrations following receptor activation. Moreover, they provide insights into the regulatory mechanisms that control signal processing by different AC isoforms.

**Author summary:** Adenylyl cyclases (ACs) are enzymes that can translate extracellular signals into the intracellular molecule cAMP, which is thus a 2^nd^ messenger of extracellular events. The brain expresses nine membrane-bound AC variants, and AC5 is the dominant form in the striatum. The striatum is the input stage of the basal ganglia, a brain structure involved in reward learning, i.e. the learning of behaviors that lead to rewarding stimuli (such as food, water, sugar, etc). During reward learning, cAMP production is crucial for strengthening the synapses from cortical neurons onto the striatal principal neurons, and its formation is dependent on several neuromodulatory systems such as dopamine and acetylcholine. It is, however, not understood how AC5 is activated by transient (subsecond) changes in the neuromodulatory signals. Here we combine several computational tools, from molecular dynamics and Brownian dynamics simulations to bioinformatics approaches, to inform and constrain a kinetic model of the AC5-dependent signaling system. We use this model to show how the specific molecular properties of AC5 can detect particular combinations of co-occuring transient changes in the neuromodulatory signals which thus result in a supralinear/synergistic cAMP production. Our results also provide insights into the computational capabilities of the different AC isoforms.

## Introduction

Information processing in the brain occurs within circuits of neurons that are interconnected via synapses. The modification of these neuronal circuits, in response to an organism’s experiences and interactions with the environment, is crucial for memory and learning, allowing the organism’s behaviour to adapt to changing conditions in its environment. One way that neuronal circuits are modified is through the process of synaptic plasticity, in which the strengths of certain synapses are either enhanced or depressed over time in response to neural activity. Insights into when plasticity happens can provide an understanding of the basic functioning of the nervous system, and its ability to learn. A very informative way to gain such insights is through analyzing the molecular circuitry of the synapses - i.e. the networks of biochemical reactions that underlie synaptic modifications. These differ across synapses, and our focus in this study is on the corticostriatal synapse, which is the interface between the cortex and the basal ganglia, a forebrain structure involved in selection of behaviour and reward learning [1,2].

All cells process information from their external and internal environment through signal transduction networks - molecular circuits evolved for producing suitable responses to different stimuli. In neurons, synaptic signal transduction networks determine whether a synapse will be potentiated or depressed. In some cases, even single molecules are able to realize computational abilities within these networks. These molecules are often enzymes, whose activity is allosterically regulated by the binding of other signaling molecules [3]. One such case is the family of mammalian adenylyl cyclase enzymes (ACs). These catalyze the conversion of adenosine triphosphate (ATP) to cyclic adenosine monophosphate (cAMP) - one of the main cellular second messenger signaling molecules.

Mammalian ACs express ten different isoforms [4–6]. Of these, nine are membrane bound, and one is soluble. Their catalytic reaction may be regulated by a variety of interactors, most importantly G protein subunits [6]. These are released in response to extracellular agonists binding to G protein-coupled receptors (GPCRs), the largest superfamily of mammalian transmembrane receptors. In this way, ACs may function not only as signal transducers but also as *signal integrators*: they perform decision functions that determine the time at which and how much cAMP is produced. One such decision function, attributed to many ACs, is detection of co-occuring signaling events (denoted as *coincidence detection* here), resulting in significantly increased production of cAMP only when more than one signaling event occurs almost simultaneously [7–15].

All nine membrane-bound AC isoforms are expressed in the brain, possibly because this organ is specialized in signal processing. Specific ACs are particularly abundant in specific brain regions, and AC5 is highly expressed in the striatum, the input nucleus of the basal ganglia. It is involved in signal transduction networks that are crucial for synaptic plasticity in the two types of medium spiny neurons (MSNs) of this brain region, which are the direct pathway MSNs that express D_1_-type dopamine receptors (D_1_ MSNs), and the indirect pathway MSNs expressing D_2_-type dopamine receptors (D_2_ MSNs).

The same regulatory mechanism exists in both MSN types: AC5 is activated by the stimulatory Gα_olf_ protein subunit, and inhibited by the inhibitory Gα_i_ protein subunit. Clearly, the regulation of AC5 is a crucial determinant of the levels of cAMP. This mechanism responds to different extracellular agonists (acting as neuromodulatory signals) associated with the expression of different G protein-coupled receptors (GPCRs) (Fig. 1A) [16–18]. In D_1_ MSNs, the binding of dopamine to the D_1_ GPCR results in the release of Gα_olf_, while binding of acetylcholine to the M_4_ GPCR causes the release of Gα_i_. Conversely, in D_2_ MSNs, binding of dopamine results in the release of Gα_i_, and Gα_olf_ is released upon binding of adenosine to the A_2a_ GPCR.

**Figure 1.**
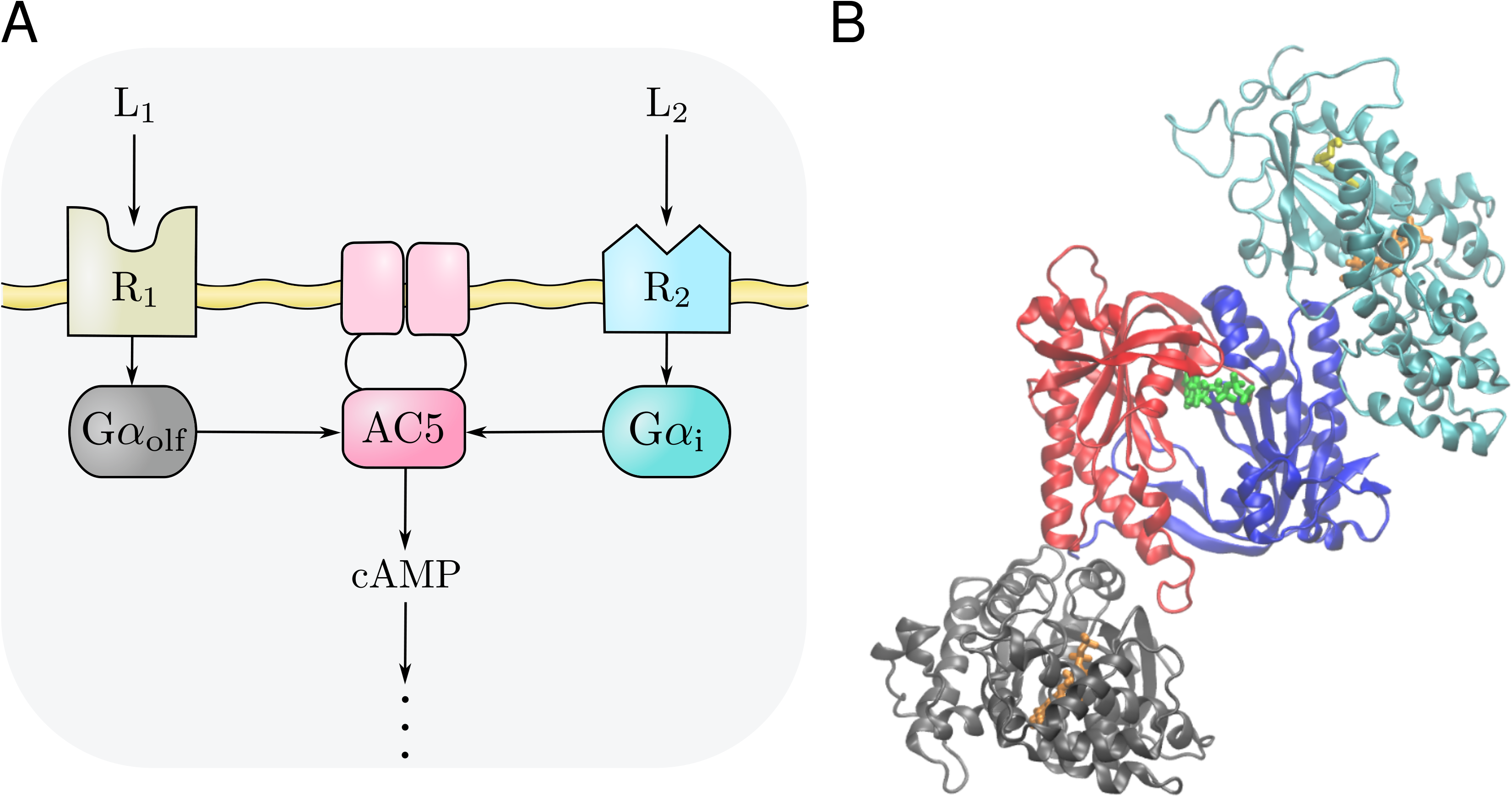
**A** General scheme of the AC5 signal transduction network. It applies to both the D_1_ and D_2_ MSNs discussed in the text. Two agonists (L1 and L2) bind to two GPCRs (R1 and R2), releasing the Gα_olf_ and Gα_i_ subunits, respectively. These stimulate and inhibit the conversion of cAMP, respectively. **B** Initial modelled configuration of the Gα_olf_ · AC5 · Gα_i_ ternary complex, used in the classical MD simulations. The cytosolic part of the AC5 enzyme consisting of the pseudo-symmetric C1 (blue) and C2 (red) cytoplasmic domains in an ATP-bound (green) conformation are complexed to an active conformation of Gα_olf_ (gray) and to Gα_i_ (cyan). GTP (orange) and the myristoyl moiety in Gα_i_ (yellow) are shown in stick representation. Controlling the relative positions and conformations of C1 and C2 may enhance or inhibit enzymatic function. This is one way in which Gα_olf_ and Gα_i_ exert their regulatory effects: each of them has a separate binding site on the AC5 domain dimer.

Knowing how AC5 processes the neuromodulatory signals would reveal the conditions under which plasticity and learning in the basal ganglia are triggered. In particular, knowing whether AC5 is a coincidence detector will help us understand if and why changes in more than one of the neuromodulatory signals it receives are necessary to trigger synaptic plasticity. This is what we have set out to determine in this study. We use the neuromodulation of D_1_ MSNs as the example here (see [15]).

Recent experimental data indicates that, during the resting state, much of AC5 in the D_1_ MSNs could be bound to Gα_i_ due to a tonic level of acetylcholine produced by the tonic activity of the striatal cholinergic interneurons [14]. Previous modeling studies of this signal transduction network predicted that AC5 responds most strongly to a simultaneous increase in dopamine (Da ↑) and a pause in acetylcholine levels (ACh ↓), i.e. both stimulation by increased Gα_olf_ and disinhibition by decreased Gα_i_ are necessary for the enzyme to produce significant amounts of cAMP [15]. The response to the two neuromodulatory signals was nonlinear and synergistic. This suggests that AC5 might function as a coincidence detector, since the network responds with significant amounts of cAMP only when the two incoming signals Da ↑ and ACh ↓ coincide in time and in the spatial vicinity of the receptors. In order to perform strong coincidence detection, the network should be able to make a clear distinction between the situation of a simultaneous dopamine peak and acetylcholine dip (Da ↑ + ACh ↓) and that of a single signal, i.e. Da ↑ or ACh ↓ alone. This distinction is realized by producing different amounts of cAMP, i.e. by differences in the enzyme’s catalytic rate.

During the course of our previous kinetic modelling study [15], and follow-up experimental studies on the function of the AC5 signal transduction network [14,19], it became clear that the presence or absence of a ternary Gα_olf_ · AC5 · Gα_i_ complex during AC5 regulation, and the level of catalytic activity of such a complex, could significantly affect AC5’s ability to act as a coincidence detector.

While the existence of the binary AC5 · Gα_i_ and AC5 · Gα_olf_ complexes, and their catalytic activities, has been confirmed experimentally, a Gα_olf_ · AC5 · Gα_i_ ternary complex has not been identified so far [20–22]. However, it has been suggested to exist during AC5 regulation, but it is not known whether it would be catalytically active or inactive (Fig. 1B) [21, 23]. So far, we know from molecular dynamics (MD) simulations of the binary complexes that binding of one Gα subunit can produce allosteric effects at the binding site for the other [24,25], raising the speculation that allosteric effects influence ternary complex formation.

Resolving the details of AC5 regulation can help to understand whether AC5 acts as a coincidence detector, and hence, whether transient changes in two neuromodulatory signals are necessary for plasticity and learning in the basal ganglia. Here, we take a multiscale modeling approach to address the following question: can the Gα_olf_ · AC5 · Gα_i_ ternary complex form in the AC5 signal transduction network? If it does, is it able to catalyse ATP conversion, and does its presence affect the ability of the enzyme to perform coincidence detection? We combine MD simulations, to study the complexation of AC5 with its Gα partners, with Brownian dynamics (BD) simulations to estimate the forward (association) rate constants of binding between AC5 and the Gα subunits. Snapshots from the MD simulations are used as starting points for BD simulations. We then incorporate the results from these simulations into a kinetic model of the AC5 signal transduction network and quantify the ability of the enzyme to detect coincident extracellular signaling events. Molecular simulation approaches that bridge multiple spatial and temporal scales are well established [26], and here we span the spatial scales by bridging from intra- and inter-molecular dynamics up to the function of a biochemical reaction network [27,28,29]. Multiscale simulation is particularly informative for this system since experimental efforts in studying the ternary complex have not yielded results.

Our MD simulations show that a ternary complex could be present during AC5 regulation, being stable on the μs timescale, but it appears to be catalytically inactive. The kinetic model, constructed using results from the MD and BD simulations performed here, reveals that: (i) the suggested interaction scheme between the regulatory Gα subunits and AC5 strengthens coincidence detection when compared to alternative schemes; (ii) the predicted values of the forward rate constants are favourable for coincidence detection. This suggests that AC5 is indeed a powerful coincidence detector and, as a result of the inactive ternary complex, a rise in dopamine alone does not have an effect on synaptic plasticity; it needs to be accompanied by a pause in the level of acetylcholine. These insights are also discussed with regard to other AC isoforms.

## Results

### Molecular dynamics simulations indicate that the apo Gα_olf_ · AC5 · Gα_i_ ternary complex is stable on microsecond timescales

The existence of a ternary complex, Gα_olf_ · AC5 · Gα_i_, during AC5 regulation has been unclear. So far it has not been possible to detect the complex in experiments, possibly due to its unstable nature [21]. However, as Gα_olf_ and Gα_i_ interact with AC5 on different sites on its catalytic domains, the ternary complex could be formed via two reactions:

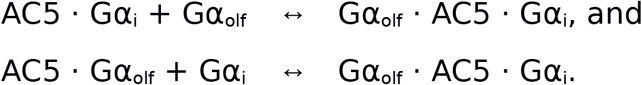

Previous MD simulation results suggested that upon binding of Gα_i_ to AC5, the Gα_olf_ binding groove adopts a conformation that hinders Gα_olf_ from binding [24], making the first reaction less favourable; however, a stable ternary complex might still be formed via the second route. Allatom MD simulations were employed to investigate the stability of a putative *apo* ternary complex, Gα_olf_ · AC5 · Gα_i_, in the absence of ATP, on the μs time scale. We mention here that Gα_olf_ has no post-translational modifications to its protein sequence. In contrast, Gα_i_ is considered in its myristoylated form since a non-myristoylated Gα_i_ subunit is known to be unable to form an AC5 · Gα_i_ complex and, therefore, it is not functional [21, 24].

The *apo* ternary complex appears to be stable over the course of 2.1 μs of all-atom MD simulation with a root-mean-square deviation (RMSD) of the complex’s backbone fluctuating between 0.8 and 1 nm (Fig. 2B) compared to the first frame of the trajectory. The RMSD of each individual protein, i.e. AC5, Gα_olf_ or Gα_i_, in the *apo* ternary complex remains below 0.4 nm (Fig. S1). Similar RMSD values have been found for the *apo* forms of the binary complexes AC5 · Gα_olf_ and AC5 · Gα_i_ (Fig. S1). Additionally, analysis of the secondary structures and calculations of the numbers of H-bonds as functions of time for all systems confirm the overall stability suggested by the RMSD analysis (Fig. S2 – S5). Root-mean-square-fluctuations (RMSF) per residue have also been calculated (Fig. S6).

**Figure 2.**
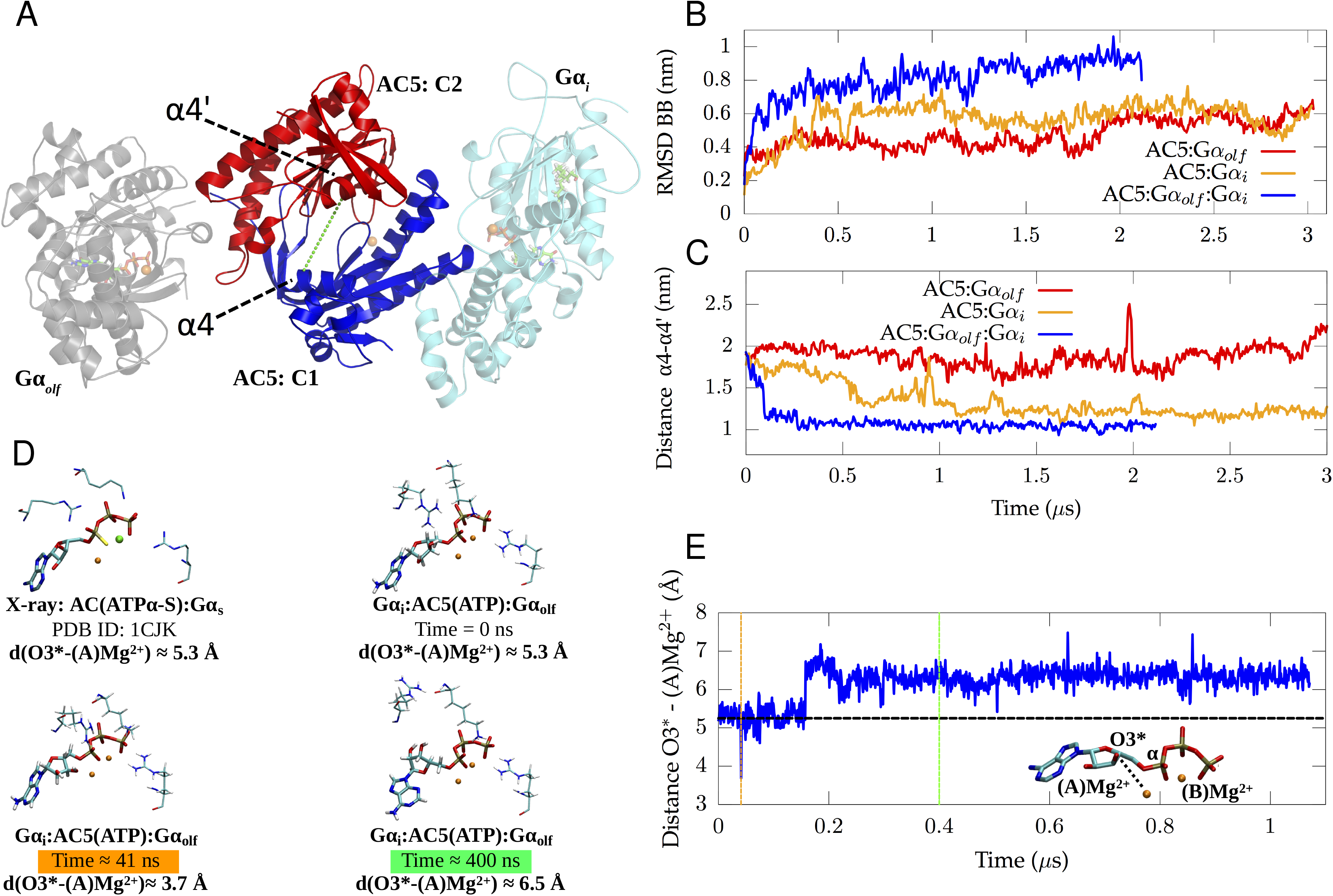
Stability analysis of the *apo* ternary complex, conformation of the active site of AC5 in the *apo* state, and conformation of ATP in AC5’s active site in the *holo* state. **A** Initial conformation of the *apo* ternary complex including Gα_olf_ (gray), Gα_i1_ (cyan), C1 (blue), C2 (red). In addition to the protein structures, two GTP molecules as well as the myristoyl moiety of the Gα_i1_ subunit and three Mg^2+^ ions are shown. The green dashed line indicates the distance between the Cα atom of Thr1007 (belonging to helix α4) and the Cα atom of Ser1208 (belonging to helix α4’). **B** RMSD of the backbone of the protein complexes Gα_olf_ · AC5 · Gα_i1_,, AC5 · Gα_olf_ and AC5 · Gα_i1_. **C** Time evolution of the distance between Thr1007 and Ser1208 for the three simulated complexes in *apo* form. **D** Conformation of ATP in the Gα_olf_ · AC5(ATP) · Gα_i1_ system at different time intervals in the trajectory as well as the conformation of ATPα-S in the AC · Gα_s_ X-ray structure (PDB code 1CJK). The color of the time in green and orange corresponds to the coloured lines in image (E). **E** Distance between the oxygen of a hydroxyl on the sugar ring of ATP, O3*, and a Mg^2+^ ion, (A) Mg^2+^, in the active site of the *holo* ternary complex. The black dashed line shows the distance between the two atoms in the AC · Gα_s_ X-ray structure (PDB ID 1CJK) to which ATPα-S is bound [22].

Minor differences in RMSF were observed for Gα_olf_ and Gα_i_, independently of the simulated system. Conversely, in the case of AC5, the C2 β4’-β5’ region (i.e. residues Val1186 to Trp1200 of the C2 domain) and the C2 β7’-β8’ region (i.e. residues Gln1235 to Asn1256 of the C2 domain) show a change in orientation dependent on the presence of the Gα subunit. Whereas β7’-β8’ appears to be more flexible in the binary AC5 · Gα_olf_ complex, the β4’-β5’ region is more flexible in the presence of Gα_i_. Besides RMSD, a measure of compactness of the simulated complexes is provided by the global radius of gyration (R_g_), which also provides an indication of the stability of the complex. R_g_ was calculated as a function of time along the simulated trajectories for all three complexes (Fig. S7). In the case of the *apo* AC5 · Gα_i_ complex, a clear reduction of R_g_ can be observed compared to its initial conformation. The *apo* form of AC5 · Gα_olf_ also shows a reduction of R_g_ over time, while R_g_ increases for the *apo* ternary complex system with respect to the starting value. In this regard, it is worth pointing out that after the first 500 ns, R_g_ does not increase further and, instead, oscillates around a constant value, slightly higher than that for the first frame. Such behaviour does not suggest instability of the ternary complex, but is rather an indication of an initial reorientation of the different domains to optimize their interaction. Such structural rearrangement is not surprising considering that the initial structure of the ternary complex was generated by homology modelling.

### Simulations of the *apo* and *holo* state of the ternary complex indicate that it is unable to catalyse ATP conversion

To assess the ability of the ternary complex to catalyse the conversion of ATP to cAMP, we investigated the structural dynamics of the ATP binding site during the MD simulation of the *apo* ternary complex, and the conformations sampled by ATP in a simulation of the *holo* ternary complex, Gα_olf_ · AC5 · Gα_i_ in which ATP is bound to the active site of AC5. We compare the conformational changes of the proteins in the *apo* ternary complex with those in the previously reported simulation of the *apo* AC5 · Gα_i_ complex [24] and with those of the *apo* AC5 · Gα_olf_ simulation reported here.

The ability of AC5 to convert ATP to cAMP depends on the state of its catalytic domain. A characteristic quantity in this respect is the relative distance between the two helices α4’ and α4 positioned either side of the binding groove (Fig. 2A). The distance between the Cα atom of Thr1007 in helix α4 and the Cα atom of Ser1208 in helix α4’ (highlighted by the green dashed line in Fig. 2A) was calculated along the simulated trajectories for the three complexes investigated here and reported as function of time in Fig. 2C.

This distance exhibits a clear decrease in the first 100 ns in the Gα_olf_ · AC5 · Gα_i_ complex, starting from a value of 1.9 nm and reaching a stable value of about 1.1 nm. Similarly, a decrease of the α4-α4’ distance was also observed in the AC5 · Gα_i_ complex along 3 μs of simulation. Conversely, when AC5 is bound to Gα_olf_ in a binary complex, the distance between the two helices is characterized by a higher value with respect to the case of the AC5 · Gα_i_ and Gα_olf_ · AC5 · Gα_i_ complexes. A larger value of this distance can be associated with a higher accessibility of the binding groove. On the other hand, a reduced value, as found in the AC5 domain bound to Gα_i_ implies a lower accessibility to the binding groove. The effect of the reduction in α4-α4’ distance on active site conformation and residue orientation has been described in [24] for the AC5 · Gα_i_ complex. A shorter distance between the α4 and α4’ helices decreases the volume in the active site accessible to ATP as α4’ starts occupying the region in which ATP’s adenine moiety docks. Due to the conformational changes in the active site, important residues for ATP stabilization undergo a rearrangement in orientation that appear to significantly differ from the AC5 · Gα_olf_ system, potentially negatively impacting the probability of ATP association. Hence, this partial closure of the active site of AC5 in AC5 · Gα_i_ and Gα_olf_ · AC5 · Gα_i_ reduces the space available for ATP binding and could also lower the probability of stable ATP association, suggesting that the ternary complex is likely to be inactive.

Apart from the ability of the *apo* ternary complex to bind the substrate, we additionally investigated the possible catalytic activity of a *holo* Gα_olf_ · AC5 · Gα_i_ complex. ATP has to undergo a cyclisation reaction in order to form the products cAMP and pyrophosphate. This reaction is induced by the attack of a deprotonated hydroxyl moiety in ATP’s sugar ring, O3*, on the phosphorus atom of the α-phosphate moiety (Fig. 2E). Hence, ATP conversion requires oxygen O3* to be in the proximity of (A)Mg^2+^ to which the phosphorus atom is coordinated in order to obtain a conformation that can undergo the cyclisation reaction [30–32].

A crystal structure of the *holo* AC domain dimer in complex with a stimulatory Gα subunit [22] shows the active conformation of ATPα-S, an ATP mimic, in the active site of the enzyme in the presence of a Mg^2+^ ion and a Mn^2+^ ion (Fig. 2D), which is assumed to be substituted by a second Mg^2+^ ion under physiological conditions. The O3*-(A)Mg^2+^ distance in the X-ray structure of the *holo* AC · Gα_s_ complex is 5.25 Å (Fig. 2D). In the MD trajectory of the *holo* ternary complex, the O3*-(A)Mg^2+^ distance starts at a similar value and mainly remains in this state for the first 160 ns. Within the first 160 ns, the O3*-(A)Mg^2+^ distance even decreases to distances as short as 3.7 Å, closer to the distance which has been suggested to correspond to ATP’s reactive state [30–32] (see orange line). However, after 160 ns, the ATP molecule undergoes a drastic conformational change, resulting in an increase of the O3*-(A)Mg^2+^ distance to ≈ 6.5 Å. In this state, the ATP molecule is unable to attain a short O3*-(A)Mg^2+^ distance, such as observed at 41 ns, which is a feature of the reactive state of ATP. This conformational transition of ATP appears to be irreversible on the μs time scale, thus suggesting the inhibition of the catalytic reaction.

### The rate of diffusional association of each Gα protein subunit to AC5 is inaffected by prior binding of the other subunit

To provide initial values for the forward rate constants - the parameters in the kinetic model of AC5 activity which were not constrained experimentally - we performed BD simulations (Table 1). The predicted rate constants suggest both Gα subunits form complexes at similar rates, and their association is not greatly affected by the presence of ATP in the AC5 active site, or prior binding of the other subunit. Indeed, the variation in the predicted rate constants for each reaction across the different MD snapshots used in BD simulations is greater than the variation of the mean values for each constant (Table S1).

**Table 1:** Bimolecular association rate constants (nM^−1^s^−1^) for the forward reactions computed from BD simulations (standard deviation over all structures of complexes and replica simulations shown in parentheses). Rate constants were calculated separately for complexes incorporating both *apo* and *holo* AC5. Data for each individual structure of each complex are shown in Table S1.

| Rate Constant | *apo* AC5 | *holo* AC5 |
| --- | --- | --- |
| $k_{f1}$ | 0.018 (0.007) | 0.026 (0.009) |
| $k_{f2}$ | 0.02 (0.01) | 0.05 (0.01) |
| $k_{f3}$ | 0.012 (0.008) | 0.017 (0.006) |
| $k_{f4}$ | 0.02 (0.01) | 0.020 (0.006) |

It should be noted that the predicted rate constants are for the diffusional approach and initial binding of the Gα subunits to AC5. The previously reported MD simulations show that the binding of Gα_olf_ to AC5 · Gα_i_ is hindered by a conformational change of its binding groove on AC5, adding an additional conformational gating contribution to the rate of binding that is not described by the BD simulation method used [24].

### The presence of an inactive ternary complex improves the ability of the network to detect coincident signals

We incorporated the results from the MD and BD simulations into a kinetic model of the AC5 signal transduction network, the basic feature of which is a regulatory scheme where the ternary complex can form (Fig. 3C). We find that this network can perform coincidence detection. To investigate how the ternary complex contributes to the network’s ability to perform coincidence detection, we compared the system with a network in which no ternary complex can form.

**Figure 3.**
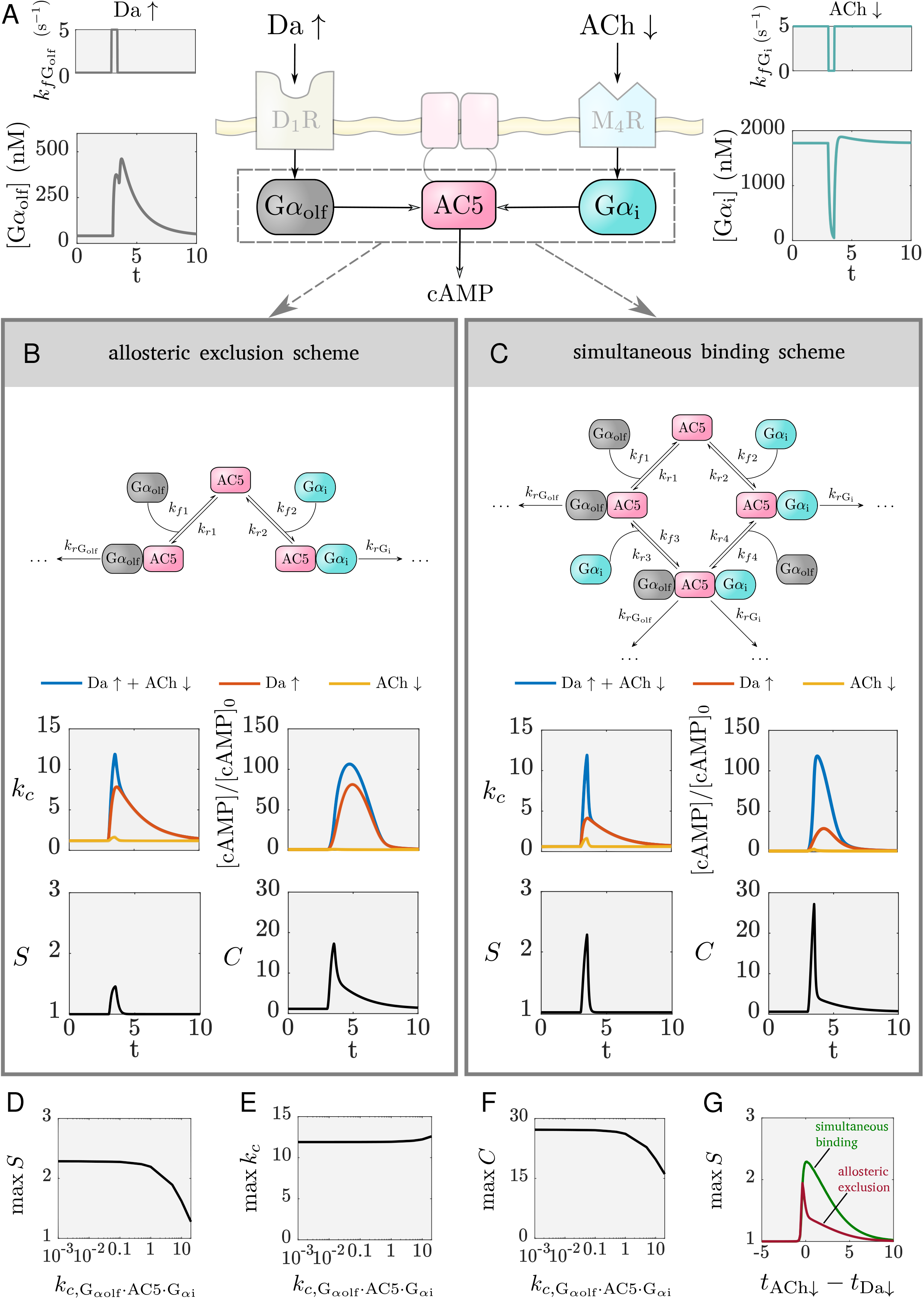
The effect of the different regulatory schemes on coincidence detection. **A** The inputs to the model translate to an elevation in Gα_olf_ and pause in Gα_i_. The shaded parts of the scheme are not included in the kinetic model. **B** The allosteric exclusion scheme, and the k_c_, synergy, and cAMP levels obtained due to this regulatory scheme. **C** The simultaneous binding scheme, and the k_c_, synergy, and cAMP levels obtained due to this regulatory scheme. **D,E,F** The effect of the ternary complex’ catalytic activity on coincidence detection: the maximum of the synergy (D), the maximum of k_c_ (E), the maximum of the metric C (F). **G** The detection window for the allosteric occlusion scheme (red) and the simultaneous binding scheme (green). t_Da_ ↑ and t_ACh_ ↓ are the times when the Da ↑ and ACh ↓, respectively. Note the shared time axes in (A), (B) and (C).

It is important to note that there are two aspects of coincidence detection: (a) distinguishing between the different inputs, and (b) responding strongly enough with an increase in cAMP concentration that is physiologically relevant. As a proxy for the amount of cAMP produced, we use the average catalytic rate of AC5, k_c_, since the amount of cAMP produced is proportional to k_c_ (see Methods, and also illustrated in Fig. 3). The average catalytic rate of AC5 is a weighted average of the catalytic rates of the unbound enzyme and each of the complexes with the Gα subunits. Additionally, to measure whether the signal transduction network distinguishes between the different input situations, the synergy quantity, S, is employed. This describes how much greater the average catalytic rate is when both signals coincide compared to the cases when only one of them arrives (see Methods). A synergy value greater than 1 indicates that the network can perform coincidence detection, i.e. the network responds more strongly when the two signals coincide than if each of them occurs individually and are summed. A value of 1, or less than 1, marks a response equal to or weaker than the sum of the individual signals, respectively. In these cases, the average catalytic rate does not produce distinguishable amounts of cAMP that enable the cell to discriminate between the different input situations. Since a synergy greater than 1 does not necessarily indicate a strong response from the network (which could happen if the basal AC5 catalytic rate is very low), a combination of the average catalytic rate and the synergy is applied where needed to quantify coincidence detection (see the metric C in methods).

The regulatory schemes that we compare are given in Figs. 3B and 3C. We name the first scheme an *allosteric exclusion scheme* – the binding of one Gα subunit excludes the possibility for binding of the other Gα subunit. The second scheme is termed the *simultaneous binding scheme*, where both Gα subunits can bind to AC5 and the ternary complex can be formed. Simulation results for both schemes are given in Fig. 3. The inputs are assumed to be a dopamine peak of 0.5s (Da ↑) and an acetylcholine dip of 0.5s (ACh ↓), and the corresponding rise in Gα_olf_ and drop in Gα_i_ are shown in Fig. 3A. Time courses showing the amounts of all AC5 species (the free enzyme and the complexes with the Gα) are given in Fig. S9.

The simultaneous binding scheme can better distinguish between Da ↑ + ACh ↓ and the individual signals Da ↑ or ACh ↓, it has higher synergy. Both the schemes have a similar maximal k_c_(Da ↑ + ACh ↓), and, as evident, the increase in synergy in the simultaneous binding scheme versus the allosteric exclusion scheme comes from a reduced k_c_(Da ↑). This relative difference in the average catalytic rate enables the simultaneous binding scheme to respond differently to the coincident signal (Da ↑ + ACh ↓) compared to Da ↑ alone, as is also visible from the amounts of cAMP produced, and hence to differentiate between the two input situations. The allosteric exclusion scheme, on the other hand, produces similar values for k_c_(Da ↑ + ACh ↓) and k_c_(Da ↑) and thus responds similarly to Da ↑ + ACh ↓ and to Da ↑ alone, being unable to distinguish well between them in terms of cAMP production. The reason for this is the exclusivity of the interaction between each of the Gα proteins and AC5: when only Da ↑ arrives, Gα_olf_ is able to compete with Gα_i_ and bind to much of the enzyme (approximately half of it as shown in Fig. S9B). This creates the catalytically active complex AC5 · Gα_olf_, driving an increase in k_c_(Da ↑). In this case, reduced inhibition of AC5 by an additional ACh ↓ does not contribute much to k_c_(Da ↑ + ACh ↓). In the simultaneous binding scheme, however, a Da ↑ causes the formation of the ternary complex (Fig. S9F), and due to its inactivity k_c_(Da ↑) does not increase significantly. Only with an additional ACh ↓ is the inhibition by Gα_i_ relieved and the proportion of the active complex AC5 · Gα_olf_ is increased, enabling a high k_c_(Da ↑ + ACh ↓). Importantly, k_c_(Da ↑) is also low for an inactive ternary complex so that little cAMP is produced with a Da ↑ only and little “stray” activation of downstream signalling would occur. In fact, only for a substantially active ternary complex does the simultaneous binding scheme become comparable to the allosteric exclusion scheme in terms of synergy (Figs. 3D, 3E, and 3F). For a wide range of low to moderate ternary complex activity, it performs coincidence detection better. The maximum of the catalytic rate is not affected much by the activity of the ternary complex (Fig. 3E), and the metric C shows that coincidence detection is most significant for an inactive ternary complex (Fig. 3F). An inactive ternary complex enables the lowest catalytic activity of the enzyme at resting state and hence the biggest difference between k_c_(Da ↑) and k_c_(Da ↑ + ACh ↓), and this in turn maximizes the synergy.

In the supplementary material we show that the allosteric exclusion scheme in itself lacks the ability to perform coincidence detection, and this is due to the exclusivity of the regulatory interaction. Coincidence detection with this scheme, as demonstrated in Fig. 3B, is in fact a result of the amounts of Gα_olf_ and Gα_i_ and the kinetics determined by the forward rate constants.

An inherent property of a coincidence detector is that there exists a time window over which two signals can be detected as if arriving together. The detector uses some mechanism by which it “remembers” the occurrence of one of the signals for some time interval, and responds when the other signal arrives within this interval. For the AC5 signal transduction network, the existence of the detection window also depends on the regulatory scheme. In fact, the formation of the ternary complex is very important to allow for a broader window of coincidence detection. In the case of only a Da ↑, a ternary complex that has buffered (or absorbed) the elevated active Gα_olf_ provides this memory: allowing the ACh ↓ to arrive some time later and still elicit a response (Fig. 3G). This is potentially relevant, since the cholinergic interneurons responsible for the ACh ↓ have been found to produce a second ACh ↓ to certain stimuli [33]. The length of the detection window is determined by the rate of deactivation of the active Gα_olf_ (also illustrated in Figs. S10D and S10G, and is due to the fact that the GTPase activity for Gα_olf_ is lower than the one for Gα_i_). Both schemes technically have the same length of detection windows, but the allosteric exclusion scheme has a high synergistic effect in a very narrow region of the window - the signals need to occur practically simultaneously (Fig. 3G). The detection window is asymmetric, i.e. the Da ↑ needs to arrive first to elicit a response from the network. (Time courses with ACh ↓ preceding and following a Da ↑ illustrating the difference between the two schemes are given in Fig. S10.)

Lastly, we note that a critical aspect for coincidence detection to work is to have fast deactivation of the active Gα_i_. Then the dynamics of Gα_i_ inside the cell can follow the short duration of the ACh ↓ signal. The experimental evidence for this high GTPase rate is listed in the description of the kinetic model (see Methods). The effect of the GTPase rate on coincidence detection is shown in Fig. S11. There is an optimum value of this rate - it needs to be both high enough to cause a drop in [Gα_i_] during the ACh ↓ and low enough so that there is significant inhibition of AC5 at steady state. Since the deactivation of Gα_i_ is faster than that for Gα_olf_, then, provided there is enough active Gα_olf_ to bind to AC5, the duration of the synergistic effect is determined by the duration of the pause (Fig. S10E and S10G). More extensive analysis on the robustness of coincidence detection to parameter values has been performed for the previous, much larger kinetic model of the signal transduction network [34].

To summarize, the results of this section show that the formation of the ternary complex aids coincidence detection and prolongs the detection window. Additionally, the less catalytically active the ternary complex is, the better the coincidence detection. As elaborated in the discussion, an inactive ternary complex can also explain recent experimental results on cAMP production due to activation of the implicated GPCRs in the native system [14,19].

### Hindrance of Gα_olf_ binding to AC5 · Gα_i_ still allows coincidence detection

To see where the predicted values from the BD simulations lie in terms of affecting coincidence detection, we performed parameter scans for the values of the forward rate constants. All of the forward rate constants affect the synergy, since they affect the fractions q_i_, i = 1, …, 4, of each species at any point in time (see equations for q_i_ in Methods).

To begin with, we address the observation from Van Keulen and Röthlisberger [24] that the Gα_olf_ binding groove in AC5 · Gα_i_ adopts a conformation that hinders Gα_olf_ binding. This could further decrease the value of the rate constant k_f4_, compared to the value predicted in the BD simulations. We investigated the effect such a decrease has on coincidence detection by altering k_f4_ by varying orders of magnitude (Figs. 4A, 4B, and 4C). We found that hindered binding of Gα_olf_ to AC5 · Gα_i_ does not affect coincidence detection significantly (the synergy S is increased by 12%). For this reason, when performing the parameter scans, we considered two scenarios. In the first one, we used k_f1_ = k_f4_ and k_f2_ = k_f3_ since the BD simulations showed that these values are similar, at least in order of magnitude. In the second scenario, we used k_f2_ = k_f3_ and k_f4_ = k_f1_/100. We call this scheme, in which the reaction corresponding to the rate k_f4_ is slower, the *hindered simultaneous binding scheme*.

**Figure 4.**
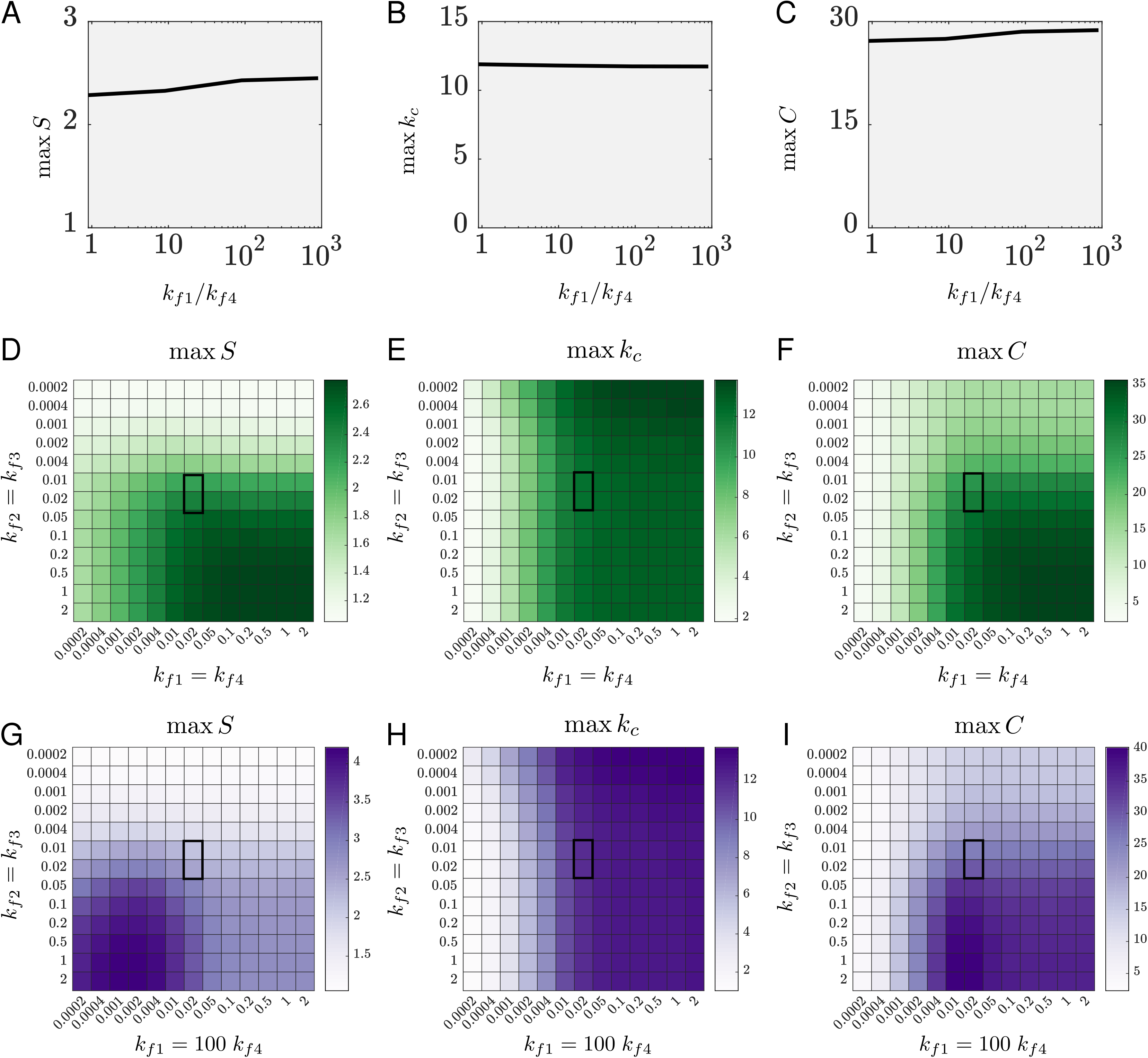
Dependence of coincidence detection on the forward rate constants. **A, B, C** The maximum of the synergy (A), the maximum of k_c_ (B) and the maximum of the metric C (C) for a reduced rate constant k_f4_ relative to the prediction from BD simulations. **D, E, F** Coincidence detection for the simultaneous binding scheme as dependent on the forward rate constants: maximum of the synergy (D), maximum of k_c_ (E), maximum of the metric C (F). **G, H, I** Coincidence detection for the hindered simultaneous binding scheme as dependent on the forward rate constants: maximum of the synergy (G), maximum of k_c_ (H), maximum of the metric C (I). Axes are in units of (nMs)^−1^. Highlighted regions correspond to the forward rate constants predicted from BD simulations.

For the simultaneous binding scheme, a very wide range of tested values for k_f1_ provides similar k_c_ and synergy values for a given k_f2_ (Figs. 4D and 4E), which can be interpreted as follows. Similarly to the requirement for a high GTPase activity for Gα_i_, it is necessary that Gα_olf_ binds to AC5 quickly enough to be able to follow the signal Da ↑. Since active Gα_olf_ has a slower dynamics than the input Da ↑, i.e. Gα_olf_ exists inside the cell for some time after the outside signal has stopped, it is likely that a range of values for the rate constant k_f1_ can enable coincidence detection (not only values that match the length of a Da ↑, which are around k_f1_ = 2 (nMs)^−1^ for a Da ↑ of 0.5s). The synergy grows with k_f1_ since a higher binding rate of Gα_olf_ provides a higher maximum of k_c_ during the Da ↑ + ACh ↓, whereas k_c_(Da ↑) is not affected much (Figs. S12A and S12B).

Increasing the rate constants k_f2_ = k_f3_ for Gα_i_ binding also causes an increase in synergy. The lower these constants, the less inhibited the enzyme will be due to smaller fractions of both AC5 · Gα_i_ and Gα_olf_ · AC5 · Gα_i_. This allows for more stimulation by the available Gα_olf_ and hence a higher resting-level k_c_ and a higher k_c_(Da ↑). The situations Da ↑ + ACh ↓ and only Da ↑ produce a more similar response, and hence show a lower synergy. Higher values for k_f2_ = k_f3_, on the other hand, allow for both stronger resting-level inhibition of AC5 and more ternary complex formed, and hence a lower resting-level k_c_ and lower k_c_(Da ↑), thus increasing the synergy. The region of optimal parameters for the simultaneous binding scheme is quite wide in terms of k_f1_ = k_f4_ and k_f2_ = k_f3_, and the most optimal scenario is for the largest values tested, as shown in Fig. 4F using the metric *C*.

Compared to the simultaneous binding scheme, the hindered simultaneous binding scheme does not affect the maximum of the average catalytic rate significantly (Fig. 4H), but it moves the region of optimal values for the synergy towards low values for the rate constant k_f1_ and increases it for these values (Fig. 4G). Inspecting the dynamics of the model with this regulatory scheme reveals the reason: the hindered reaction AC5 · Gα_i_ + Gα_olf_ ↔ Gα_olf_ · AC5 · Gα_i_ causes less ternary complex to be formed and consequently less AC5 · Gα_olf_ from the dissociation of Gα_i_ from the ternary complex (the route via k_r3_) (Figs. S12C and S12D). This, in general, lowers k_c_(Da ↑) compared to the simultaneous binding scheme, creating a larger relative difference between k_c_(Da ↑) and k_c_(Da ↑ + ACh ↓) which results in a high synergy. This difference is largest for low values of k_f1_ and decreases with k_f1_ due to the higher amounts of AC5 · Gα_olf_ during the Da ↑ (higher k_c_(Da ↑)). Importantly, the maximal k_c_ has the opposite dependence on k_f1_ from the synergy, so that the region of values for k_f1_ optimal for coincidence detection optimizes between a high catalytic rate and a high synergy (Fig. 4I). The effect of k_f2_ = k_f3_ on coincidence detection is the same as described for the simultaneous binding scheme.

In both scenarios, the predictions of the BD simulations are favourable for coincidence detection. However, higher binding rates of Gα_i_ than the predicted ones are better suited for coincidence detection. This increase in k_f2_ and k_f3_ could arise if Gα_i_ were part of multiprotein signaling complexes, as has been shown for AC5 and Gα_olf_ [35–37].

## Discussion

In this study we find that an inactive ternary complex between AC5 and its G protein regulators, a molecular-level feature, gives rise to significantly increased coincidence detection, a systems-level function of the signal transduction network that AC5 is embedded in. In order to investigate the stability and the activity of the putative Gα_olf_ · AC5 · Gα_i_ ternary complex, we carried out all-atom MD simulations. Our results showed that on the μs time scale, the complex seems to be stable independently of the presence or absence of ATP. Additionally, previous MD studies suggest possible pathways for the formation of the ternary complex, showing the possibility of the Gα_i_ protein to bind to the *holo* AC5 · Gα_s_ complex [25], and disfavoring the binding of Gα_olf_ to the *apo* AC5 · Gα_i_ complex [24]. Overall, it should be pointed out that MD simulations cannot exclude the instability of the ternary complex on longer time scales, which cannot be assessed due to computational limits. However, MD simulations can help us to investigate the conformations sampled by the ternary complex at physiological temperature and pressure and, consequently, the activity of the complex. Indeed, the partial closure of the binding groove found in the *apo* ternary complex, analogous to that occurring in the binary *apo* AC5 · Gα_i_, suggests a lack of catalytic activity in the ternary complex due to the reduced accessibility for the ATP substrate. Additionally, even if a ternary complex could exist with ATP bound to AC5, our results show that the substrate would adopt a conformation not suitable for the subsequent catalytic reaction leading to the formation of cAMP. It is worth mentioning that possible conformations sampled by ATP, prior and during its conversion to cAMP, have been reported by Hahn et al. in a theoretical study [38]. Such conformations, required for an optimal conversion of ATP to cAMP, clearly show an opposite orientation of the oxygen O3* with respect to that sampled in our simulation, thus supporting our hypothesis about the inactivity of the ternary complex.

Experimentally, indirect data exist that are consistent with a significantly inactived ternary complex. To begin with, the existence of functional oligomeric complexes of AC5 and the A_2a_ and D_2_ receptors has been demonstrated, and their proposed spatial arrangement, importantly, supports ternary complex formation [39]. Additionally, experimental results from striatal cultures in this study show that stimulating the D_2_ receptor almost entirely counteracts the effects of the applied agonist on the A_2a_ receptor in terms of cAMP production. These results are consistent with an inactive ternary complex since, under these conditions, both active Gα_olf_ and active Gα_i_ would exist in the cell, and activation of Gα_i_ stops the activity of the previously formed AC5 · Gα_olf_. Similar experimental studies with the *in vitro* native system have also been performed recently [14,19], although the animals used in these studies are very young (between 7 and 12 days old) and the developmental transition from AC1 to AC5 in the striatum is not complete. However, at P7 AC5 contributes to at least 50% of the cAMP response, and this contribution significantly increases further on, as the transition is reported to be complete by P14 [40]. Comparison with these experimental results is therefore still informative, and in both studies the results are consistent with an inactive ternary complex. In D_1_ MSNs, stimulating the D_1_ receptors with an agonist followed by activation of the M_4_ receptors completely abolishes the cAMP response of AC5 [14, Fig. 3A]. The analogous experiment in D_2_ MSNs is also consistent with an existing and inactive ternary complex [19, Fig. 2B]. Stimulating the A_2a_ receptor with an agonist and then uncaging dopamine, which stimulates the D_2_ receptor, also abolishes the cAMP response. Experiments modeling the opposite regulatory effects of Gα_s_ and Gα_i_ on AC5 in living HEK293T cells via two other GPCRs are also consistent with an inactive ternary complex, where activation of the Gα_i_ signaling branch completely reverts cAMP production due to the activated Gα_s_ branch back to baseline levels [41, Figs. 6A,B]. Earlier *in vitro* experiments with membranes of Sf9 cells expressing AC5, however, have not yielded complete inhibition by Gα_i_ [23,42]. Adding high amounts of both Gα_olf_ and Gα_i_ to the assays of AC5-containing membranes did not completely inhibit AC5 - the enzyme still produced significant amounts of cAMP [23,42]. Nevertheless, a catalytically active ternary complex still supports coincidence detection for a wide range of ternary complex activity, as shown in Figs. 3D-F, as long as its catalytic rate is noticeably smaller than that of the active binary complex AC5 · Gα_olf_. Additionally, the data in [23] are interpreted and fitted with an alternative, more elaborate interaction scheme where the ternary complex can form and is not very active, and the production of cAMP is due to higher order, catalytically active complexes of AC5 with more than one Gα_olf_ and Gα_i_ subunit. We therefore suggest that the results of our study are plausible and supported by several experimental results, among which are experiments with the native system.

### Relevance for corticostriatal synaptic plasticity

Knowing what an intracellular signal transduction network is composed of and the details of how it operates can help to clarify how it responds to extracellular events. The AC5 signal transduction network in D_1_ MSNs (as well as in D_2_ MSNs) interacts with a calcium-activated signal transduction network to regulate synaptic plasticity. Calcium influx at the synapse is necessary for synaptic potentiation, but exerts its effect only if accompanied activation of the AC5 signaling module [43–45]. Thus, the AC5 signal transduction network gates plasticity in the corticostriatal synapses onto the MSNs. Now, an existing and inactive ternary complex in AC5 regulation has consequences on how this “gate” would be opened: disinhibition from active Gα_i_ is necessary, accompanied by stimulation from Gα_olf_. That is, our findings suggest that experimentally observed ACh ↓ in the striatum is likely physiologically relevant for D_1_ MSNs, and both a Da ↑ and a ACh ↓ are necessary to enable synaptic potentiation (see also [46]). They can also help in interpreting the functional role of the neuromodulatory signals in the striatum.

The kinetic model of the AC5 signal transduction network, built according to the findings of the MD simulations and with the parameters predicted with the BD simulations, reveals improved coincidence detection when compared to alternative regulatory schemes. This, together with the implications mentioned above arising from the existence and inactivity of the ternary complex, suggests that the regulation of AC5 has indeed evolved to perform coincidence detection of the two neuromodulatory signals.

### Comparisons to other AC isoforms and AC-dependent cascades

In this study we have investigated the regulation of AC5 through interaction with the Gα_olf_ and Gα_i_ subunits. All membrane-bound AC isoforms are known to be stimulated by Gα_s_, a close homologue of Gα_olf_, while only ACs 1, 5 and 6 are inhibited by Gα_i_ [6]. With this in mind, it is interesting to ask whether our findings concerning AC5 regulation, particularly the presence of an inactive ternary complex in the signalling network, could also be valid for cascades containing ACs 1 and 6. The sequences of rat Gα_s_ and Gα_olf_ are highly similar with an identity of 76.0 % and similarity of 90.0 % (Fig. S13). Restricting the comparison to the amino acid residues within 6 Å of AC5 in the modelled *apo* AC5 · Gα_olf_ complex, the identity rises to 95.8 %, with only a single position differing. For this reason, it is reasonable to assume that our findings regarding the formation of Gα_olf_-containing complexes are also applicable to Gα_s_. Indeed, our modelling of the AC5 · Gα_olf_ complexes assumes that we can take crystal structures of AC · Gα_s_ complexes as template structures. It should be noted that Gα_s_ is known to be deactivated more slowly than Gα_olf_ [47], which could reduce the ability of Gα_s_-activated networks to detect subsecond coincident signals (see [15]).

Previously, we have performed electrostatic analyses of mouse AC isoforms, to identify regions of electrostatic similarity within similarly regulated isoforms [48]. In that work, we showed that the molecular electrostatic potentials surrounding ACs 1, 5 and 6 in aqueous solution were very similar in the Gα_i_ binding region of AC5, suggesting that the location of binding, and the bound orientation, of Gα_i_ on ACs 1 and 6, could be similar. This electrostatic similarity was due to a more negative character, compared to other AC isoforms, which is complementary to the largely positive potential of the face of Gα_i_ that contains the switch II region that is thought to interact with AC.

The sequence identities of AC isoforms with respect to AC5, in the C1 domain to which Gα_i_ binds, show that ACs 1 and 6 are most similar, at 73.5 % and 94.7 %, respectively (Fig. 5B). Considering only those residues predicted to be involved in the interaction with Gα_i_, the similarities are 64.7 % and 91.2 %. From the very high electrostatic and sequence similarity, it seems reasonable to assume that our findings should hold for AC6.

**Figure 5.**
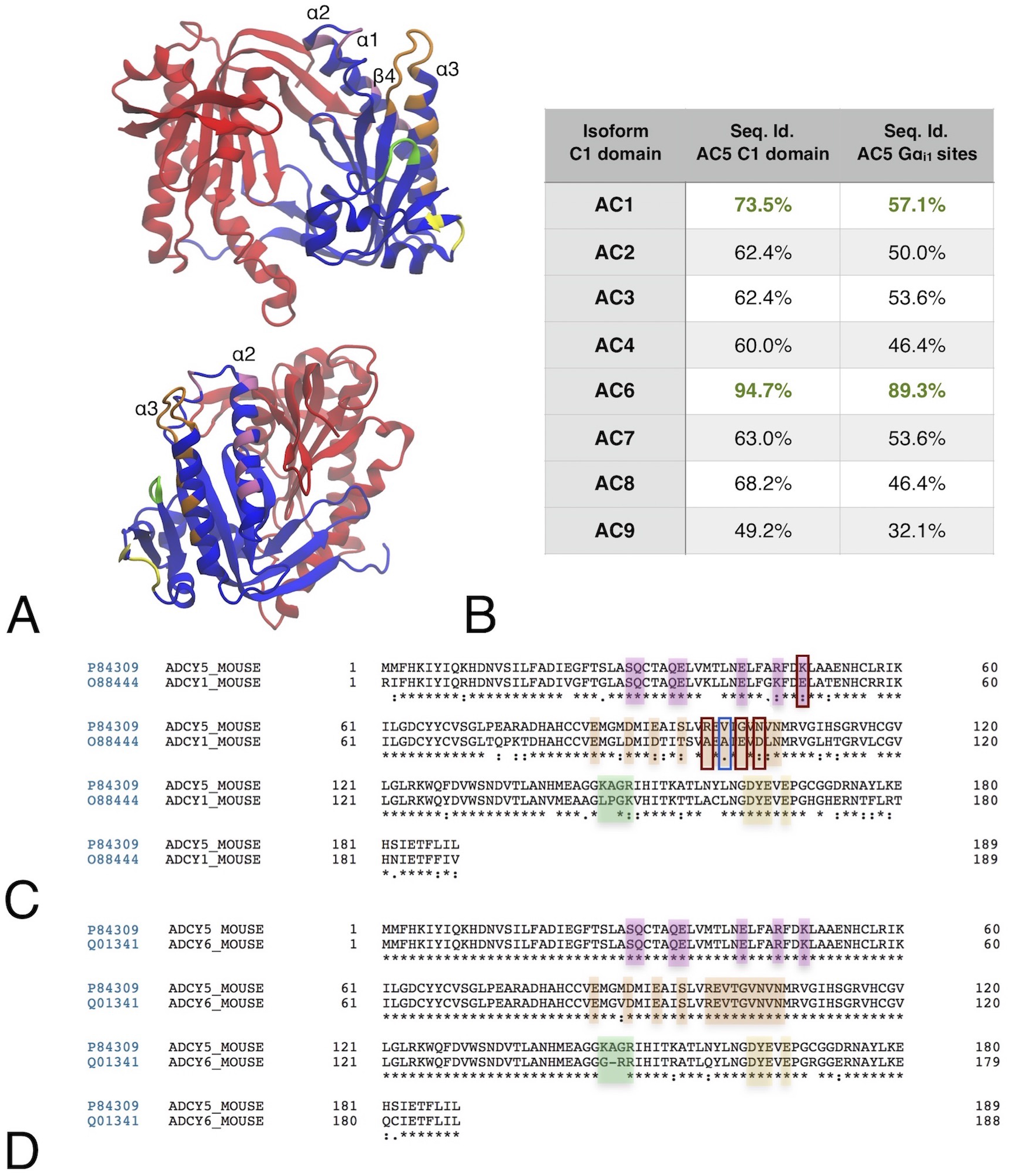
**A** The structure of the C1 (blue and highlight colors) and C2 (red) catalytic domains of AC5. **B** The highlighted regions show the AC5 residues that interact with Gα_i_ during complex formation. Overall sequence similarity for the C1 domain for the other mouse AC isoforms, compared to mouse AC5, and for the highlighted residues that interact with Gα_i_. **C, D** Pairwise sequence alignments for AC1 (C) and AC6 (D) with the colors matching those of the structure in subfigure A. The mouse sequences were taken from Uniprot, and aligned using Clustal Omega within Uniprot. The C1 domain was identified through alignment with PDB 1CJK [22], which contains a canine AC5 C1 domain, and the residues are numbered from the start of the C1 domain. The first residues of the C1 domains are at positions 288, 456 and 363 in the canonical sequences of AC1, AC5 and AC6 respectively. The red boxes in (C) indicate charge-altering substitutions between the AC5 and AC1 sequences in the Gα_i_ binding α3 helix, while the blue box indicates a substitution in a residue whose mutation was found to affect Gα_i_-induced inhibition of AC5 [21].

While AC1 is inhibited by Gα_i_, the level of inhibition is much lower, with higher levels only seen for its forskolin or calmodulin-activated states [49], therefore directly applying our findings to AC1 cascades is more difficult. In the AC1 C1 domain sequence, there are three charge-altering substitutions in the region formed by the C-terminal end of α3-helix and the loop connecting this helix to the β4-strand (Fig. 5C). These substitutions give this region a more negative character, and therefore it is reasonable to assume that they should not destabilize the binding mode we find for Gα_i_ on AC5 to a large extent. In a previous mutagenesis study [21], including one of these substitutions (N559D by Uniprot sequence, N480D by sequence in [21]), which is also present in the non-Gα_i_ inhibited ACs 2, 4, 7 and 8 (Fig. S14), as a mutation in AC5 was found to produce only a small reduction in its inhibition by Gα_i_ (less than 2-fold increase in IC50). A more significant effect on the binding of Gα_i_ to AC1 may occur due to a difference in the C-terminal residue of the α3-helix of the C1 domain. In AC1, there is an alanine in this position, while in ACs 5 and 6, it is a valine. The wildtype AC5 construct used in [21] differed from the canonical sequence by having a methionine in this position (476 by their sequence numbering, 555 in the Uniprot sequence). They showed that mutating this residue to match the canonical sequence reduced the IC50 of Gα_i_ by a third, while mutation to alanine gave a greater than 30-fold increase in IC50. Due to this apparent reduction in the affinity of AC1 for Gα_i_, further MD simulations may be required to confirm the stability of a Gα_olf_ · AC1 · Gα_i_ ternary complex. The lower sequence similarity between the C1 domains of ACs 1 and 5 also suggests that the allosteric effects on both the Gα_olf_/Gα_s_ binding groove and the active site could be different in a putative AC1 ternary complex. Again further MD simulations would be required to investigate this, as well as to further unravel the different computational properties of the AC isoforms found in different synapses.

### Assumptions and limitations

As for any simulation study aimed at investigating *in vivo* subcellular processes, there are limitations that should be discussed. First, adenylyl cyclases, together with other components of the signal transduction networks they participate in, are organized as parts of multiprotein signaling complexes and/or are localized in structured microdomains in the cell which serve to compartmentalize the effects of the produced cAMP and efficiently activate downstream components of the networks and, ultimately, effectors [35–37, 39, 50]. The kinetic model here, on the other hand, assumes mass-action kinetics for the species included, i.e. it disregards any organization into multiprotein signaling complexes and instead describes a well-mixed solution of molecular species. This means that it reproduces the experimental measurements on cAMP in a partly phenomenological way. For example, AC5 needs to be presented with appropriate proportions of Gα_olf_ and Gα_i_ to reproduce the levels of cAMP, which may not necessarily be the same as the amounts of these proteins in real synapses. Second, a recent study has demonstrated that AC5 and the heterotrimeric G_olf_ protein are pre-assembled into a signaling complex and suggested that upon activation by the GPCR, the Gα_olf_ subunit rearranges rather than physically dissociates from the Gβγ subunit [37]. This would imply an increase in the forward rate constant for AC5 and Gα_olf_ binding predicted with the BD simulations. In light of this study, the effect of the hindered Gα_olf_ binding to AC5 · Gα_i_ is not clear, since its main advantage of improving coincidence detection occurs precisely for values of k_f1_ around the BD estimates. It remains to be seen how much of an increase in the forward rate constant preassembly confers. A similar situation likely occurs for Gα_i_, as well - the GPCRs and AC5 form oligomeric complexes that could include the G proteins [39], which would mean that the predicted value for the forward rate constant for AC5 and Gα_i_ binding, k_f2_, is also probably an underestimate. A larger value for k_f2_ is beneficial for coincidence detection both with the simultaneous and the hindered simultaneous binding scheme (Figs. 4G–I). Finally, we should underline that the enzyme is additionally regulated by protein kinase A (PKA), calcium ions, nitric oxide, and the Gβγ subunit, and regulation via the transmembrane domains has been proposed (but not demonstrated so far) [4, 35, 51]. PKA is activated by cAMP and is the most common kinase to elicit the various downstream responses of the signal transduction network. PKA also inhibits AC5 via phosphorylation, and this is probably feedback that serves for signal termination. Calcium also inhibits AC5, an interaction which, due to excitatory synaptic input, might also help terminate its activity. The Gβγ enhances the effect of Gα_s_ on AC5, but has no effect alone, and nitric oxide has an inhibitory effect whose purpose is also unknown. The measures for coincidence detection used here do not include these additional regulatory interactions, and this would provide a more complete picture of the enzyme’s regulation in studies of various signaling scenarios.

On the same topic of molecular organization, while it is known that the D_2_ and A_2a_ receptors are colocalized and form oligomeric complexes in D_2_ MSNs, the same has not been demonstrated for the D_1_ and M_4_ receptors in D_1_ MSNs. However, the D_1_ and M_4_ receptors have individually been reported in the postsynaptic density in striatal tissue (among other compartments) by electron microscopy [52,53], and activation of the M_4_ receptors counteracts activation of the D_1_ receptors both in striatal slices, as described above, and in striatal membrane preparations [17]. This makes it likely that they indeed colocalize enough to enable both G proteins to interact with AC5.

Furthermore, concerning the structural simulations carried out in the present study, it is appropriate to highlight some aspects. First, while the *apo* and the *holo* ternary complexes are relatively stable over about 2.1 and 1.1 μs of MD simulation time, respectively, we cannot exclude that on a longer time scale such complexes could show a dissociation of the different protein units. In this regard, it is worth mentioning that both simulations of the *apo* and *holo* forms of the ternary complex have been repeated starting with new velocities for about 0.6 and 1.0 μs, respectively, leading to the same overall conclusions described above. Second, in the systems simulated here, only the catalytic domains of AC5 are considered, while the transmembrane domains are not included. Although the transmembrane domains are important for the proper dimerization of the catalytic domains [54], the functionality of AC5 was experimentally found to be maintained upon removal of the transmembrane domains [21, 55]. It should also be noted that the structure of the AC5 catalytic domain modelled in this study does not include the C1b domain (whose structure has not yet been determined). The absence of this domain has been shown to reduce the sensitivity of AC5 to Gα_i_ [21]. Nevertheless, the MD simulations performed here and by Van Keulen and Röthlisberger [24] show that Gα_i_ is able to inhibit the activity of AC5 in the absence of the C1b domain. This suggests that the lower inhibition is due to the reduced affinity of Gα_i_ in the absence of C1b, rather than do to a change in the internal mechanism of inhibition. Were it possible to include C1b in the BD simulations, this might lead to an increase in the rate of Gα_i_ association, which would lead to increased synergy in both the simultaneous and hindered binding schemes.

As mentioned above, the membrane-anchoring is also neglected in the BD simulations in which freely diffusing solutes are assumed. This assumption has two main effects on the predicted association rate constants, which alter the predictions in opposite directions, leading to some degree of cancellation of errors. The anchoring of AC5 and Gα_olf_ in the membrane would add additional diffusional constraints that would potentially increase rates in vivo, by reducing the search space, while the slower diffusion of the lipid anchor in the membrane would slow down association.

It should also be noted that the simulated systems in this work were built by homology modelling using the then available X-ray crystal structures of AC as described in the Methods section and also reported in previous studies [24, 56]. While this paper was in review, structures of AC9 including its two six-helix transmembrane domains (but not the C1b domain) were reported [57]. In order to check possible overlaps between atoms of the Gα_i_ protein in our modelled ternary complex and atoms of the new AC9 structure (and of the membrane bilayer), we fitted the Cα atoms of the C1a and C2a domains on the respective atoms of the AC9 protein (see Fig. S15). The figure clearly shows that the position of the Gα_i_ protein in the ternary complex does not overlap with the membrane. Moreover, the conformation, orientation and binding mode of the Gα_i_ to AC5 would also be perfectly suitable for AC9 complexation, providing additional support for our model. Finally, although not all parts of the AC9 enzyme could be resolved, this new cryo-EM structure could provide a basis for future modelling work to investigate the effects of membrane tethering on AC5 regulation.

## Conclusions

In this work, we have investigated the stability and activity of a Gα_olf_ · AC5 · Gα_i_ ternary complex by MD simulation and found that the complex appears to be stable on the microsecond time scale, but is unable to catalyze ATP conversion. Using BD simulations, we have made predictions of the association rate constants for the formation of both binary AC5 · Gα complexes, and the subsequent association of the second Gα subunit to form the ternary complex.

Kinetic modelling of the AC5 signal transduction network showed that the predictions of the structure-based simulations maximize the ability of the network to recognize coincident neuromodulatory signals, with coincidence detection enhanced by both the presence of the ternary complex, and its reduced activity. Additionally, coincidence detection is enhanced by the hindered binding of Gα_olf_ to the the binary AC5 · Gα_i_ complex, as suggested by our previous MD study [24]. Taken together, these results provide evidence that AC5 has evolved to perform coincidence detection of transient changes in the amounts of Gα_olf_ and Gα_i_ proteins, such that a brief deactivation of the Gα_i_ signaling branch is needed to gate the Gα_olf_ signal through. For the corticostriatal synapse on D_1_ MSNs, this implies that both the transient rise in dopamine and the decrease in acetylcholine levels are necessary to trigger synaptic plasticity.

## Methods

### Modelling of Binary AC5 Complexes

The crystal structure (PDB ID 1AZS) of the ATP-free AC · Gα_s_ complex [20] was used as a template for the catalytic region of AC5 and Gα_olf_ in the *apo* ternary complex. The template used for the initial complex conformation included 1AZS’s C1 and C2 domains (more specifically, canine AC5 for C1 and rat AC2 for C2) for modelling the AC5 structure (UniprotKB Q04400) from *Rattus norvegicus* as well as the Gα_s_’ structure for the initial Gα_olf_ (UniprotKB P38406) conformation from *Rattus norvegicus* [20, 58, 59]. The modelled structure of the myristoylated *Rattus norvegicus* Gα_i_ subunit (UniprotKB P10824) interacting with guanosine-5’-triphosphate (GTP) and Mg^2+^ was taken from [56]. Gαi also has several isoforms, and the one referred to here is Gαi1. Myristoylation is a post-translational modification of the N-terminus of Gα_i_ that results in the covalent attachment of a 14-carbon saturated fatty acid to the N-terminal glycine residue of Gα_i_ via an amide bond. The modelled AC5 and Gα_i_ structures were used for docking Gα_i_ on AC5’s C1 domain to finalise the initial conformation of the ternary complex. This *apo* ternary complex setup was also used to simulate the *apo* forms of AC5 · Gα_i_ and AC5 · Gα_olf_ by removing the subunit not to be considered.

The crystal structure (PDB ID 1CJK) of the catalytic AC domains with a bound ATP analogue (Adenosine-5’-rp-alpha-thio-triphosphate), ATPαS, and a Gα_s_ interacting with the AC protein, was used as a template for the *holo* ternary complex. The template used for modelling the active AC5 (UniprotKB Q04400) conformation in the ternary complex included the C1 and C2 domains from the PDB file 1CJK [22, 58, 59]. The Gα_s_ subunit present in 1CJK was used as a template for the initial Gα_olf_ (P38406) structure from *Rattus norvegicus* [22, 58, 59]. The modelled structure of the myristoylated *Rattus norvegicus* Gα_i_ subunit (UniprotKB P10824) interacting with GTP and Mg^2+^ was taken from [56]. The active myristoylated *Rattus norvegicus* Gα_i_ is referred to simply as Gα_i_ because only a myristoylated form of Gα_i_ was used in all simulations.

### Modelling of Ternary AC5 Complexes

Membrane-bound ACs consist of two membrane-bound regions and two cytosolic domains. The latter form the active site of the enzyme and its structure has been determined by crystallography. The catalytic domains, C1 and C2, are located close to the membrane due to AC5’s transmembrane domains, but remain entirely solvated. The crystal structure templates, used for modelling the complexes, were employed to determine the orientation of Gα_olf_ on the C2 domain. The HADDOCK web server [60] was used for docking ten conformations of the active Gα_i_ subunit to AC5’s catalytic domains in the *apo* and *holo* forms as described in [24]. The active region of Gα_i_ was defined in HADDOCK as a large part of the alpha helical domain (112-167), the switch I region (175-189) and the switch II region (200-220), allowing for a large unbiased area on the Gα_i_ protein surface to be taken into account during docking. The active region of AC5’s C1 domain was defined as the α1 helix (479-490) and the C-terminal region of the α3 helix (554-561) because experimentally it has been found that Gα_i_ is unable to interact with C2 and its main interactions with AC5 are with the C1 domain [21]. Ten snapshots of Gα_i_ were used for docking the Gα subunit to the catalytic domain of AC5. These snapshots were extracted at time intervals of 0.5 ns from the end of the classical MD trajectory of Gα_i_ (around 1.9 μs) described in [56]. The same three criteria for complex selection as in [24] were applied: (1) the absence of overlap between the C2 domain and Gα_i_, (2) no overlap with the GTP binding region of Gα_i_ and the C1 domain of AC5, and (3) presence of similar complexes in the top-ten docking results of the docking calculations performed for all ten used Gα_i_ conformations.

### Classical Molecular Dynamics Simulations

The Gromacs 5.1.2 software [61, 62] was used to perform the simulations. The *apo* and *holo* Gα_olf_ · AC5 · Gα_i_ systems, which were simulated for 2.1 μs and 1.2 μs respectively, each include two GTP molecules. In addition, the *holo* complex incorporates four Mg^2+^ ions and one ATP molecule, while in the *apo* form only three Mg^2+^ ions are present. Both *apo* and the *holo* systems were solvated in about 162,000 water molecules and 150 mM KCl. They were simulated at a temperature of 310 K and a pressure of 1 bar, maintained using the Nosé-Hoover thermostat and an isotropic Parrinello-Rahman barostat, respectively. The force fields used for the protein and the water molecules were AMBER99SB [63] and TIP3P [64]. For GTP and ATP, the force field generated by Meagher et al. was used [65]. The adjusted force field parameters for Cl^−^ and K^+^ were taken from [66]. The Mg^2+^ parameters originated from [67] and the parameter set for the myristoyl group was taken from [56]. Electrostatic interactions were calculated with the Ewald particle mesh method with a real space cutoff of 12Å. The van der Waals interactions also had a cutoff of 12 Å. Bonds involving hydrogen atoms were constrained using the LINCS algorithm [68] The integration time step was set to 2 fs.

The *apo* binary complexes, AC5 · Gα_i_ and AC5 · Gα_olf_, used for comparison to the ternary complexes, were built via the same procedure and were each simulated for 3 μs.

The first step in the equilibration procedure of the protein systems included the energy minimisation of the protein complex together with the ligands (Mg^2+^, GTP, ATP), referred to as the complete complex. Position restraints of 1000 kJ mol^−1^ nm^−2^ were applied to the structure of the complete complex during energy minimisation. The next step that was performed was the simulation of the complete complex under canonical NVT (constant number of atoms (N), constant volume (V) and constant temperature (T)) conditions, starting from the energy minimised structure, with position restraints of 1000 kJ mol^−1^ nm^−2^ on the complete complex. The length of the NVT run was 2 ns. The third step included the performance of a 4 ns isothermal-isobaric NPT run (constant number of atoms (N), constant pressure (P) and constant temperature (T)) with position restraints of 1000 kJ mol^−1^ nm^−2^ on the complete complex. The fourth step was an NPT simulation of 4 ns, with position restraints of 1000 kJ mol^−1^ nm^−2^ on the complete complex except for the hydrogens of the proteins. The fifth step contained an NPT run of 4 ns in which the backbone of the proteins were restrained by 1000 kJ mol^−1^ nm^−2^ as well as the ligands. The sixth step was an unrestrained NPT simulation of at least 10 ns, which was prolonged depending on the RMSD convergence of the proteins in the equilibrated system.

### Forward Rate Constant Estimation via Brownian Dynamics Simulations

Brownian dynamics (BD) simulations were performed to estimate the forward association rate constants in the two schemes in Fig. 3. The simulations were performed using the SDA 7 software package [69], with the associating species represented in atomic resolution as rigid bodies. The inter-species interactions were modeled using the effective charge model [70], with the electrostatic desolvation term described by Elcock et al. [71], and following the parameterization of Gabdoulline and Wade [72].

The atomic structures of the reactant species in the forward reactions shown in Figs. 3B and 3C were taken from clustered snapshots of the MD simulations described above, except for the structures of *apo* AC5 and Gα_i_ used to calculate k_f2_, which were obtained from simulations performed by Van Keulen and Röthlisberger [24]. The electrostatic potential of each snapshot of the reactant species was calculated via solution of the linearized Poisson-Boltzmann equation (PBE) using the APBS PBE solver [73], such that at the grid boundaries, the electrostatic potentials matched those of atom-centered Debye-Hückel spheres. The atomic charges of the protein residues were taken from the Amber force field, with the charges of the myristoyl moiety as described in [56] and the GTP charges as described by Meagher et al. [65]. The low-dielectric cavity (ε_r_ = 4) was described using Bondi atomic radii [74] and the smoothed molecular surface definition of Bruccoleri et al. [75], while the solvent was modelled using a dielectric constant of 78, and a 150 mM concentration of salt with monovalent ions of radius 1.5 Å. For the single species reactants, solution of the linearized PBE generated cubic potential grids with 129 grid points per dimension, spaced at 1 Å, while larger grids with 161 points per dimension were generated for the binary reactants. Effective charges were calculated using the *ECM* module of SDA 7 [69, 70], with charge sites placed on the heteroatoms of ionized amino acid side chains and termini, the phosphate oxygen and phosphorus atoms of ATP and GTP, and Mg^2+^ ions. The effective charges of each solute were fitted such that, in a homogeneous dielectric environment, they reproduced the solute’s electrostatic potential computed by solving the PBE in a skin bound by the surfaces described by rolling probe spheres of radii 4 and 7 Å along the molecular surface of the solute. Electrostatic desolvation potentials were calculated using the *make_edhdlj_grid* tool in SDA 7 [69].

For each reaction, four BD simulations of 50 000 trajectories were performed using each MD snapshot, and rate constants were calculated using the Northrup, Allison and McCammon formalism [76]. The mean and standard deviations across these four simulations was then determined. The average value for the corresponding rate constant was computed across all MD snapshots. The infinite dilution diffusion coefficients of each reaction species were calculated using HYDROPRO [77, 78] with the exception of those of the AC5 · Gα complex reactants in the ternary complex forming reactions, for which the diffusion coefficients of AC5 were used. In each simulation trajectory, the position of the center of AC5, or the reactant complex, was fixed at the center of the simulated volume, while the initial position of the center of the reactant Gα subunit was placed on the surface of a sphere of radius *b*, centered on the other reactant, with *b* taken as equal to the sum of the maximal extent of the distance of an atom of either reactant from the reactant center, plus the maximal extent of any interaction grid point to the solute center plus 30 Å. The simulations continued until the reactants diffused to a separation *c*, where *c =* 3*b*. Trajectories were assumed to have formed reactive encounter complexes when two independent native contacts between the two reactants reach a separation of 6.5 Å or less. Native contacts were defined as a pair of hydrogen bonding heteroatoms, separated by less than 5 Å in the bound complex. Two native contacts were assumed to be independent if the heteroatoms on the same solute that form the contacts were separated by more than 6 Å. This definition of an encounter complex has been shown to result in calculated proteinprotein association rate constants that correlate well with experimental values [72].

### Kinetic Model of the Signal Transduction Network

The kinetic model of the signal transduction network is a system of coupled ordinary differential equations with mass-action kinetics modeling the network’s biochemical reactions. For example, for the reaction 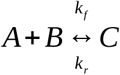, the rate at which it occurs is given with:

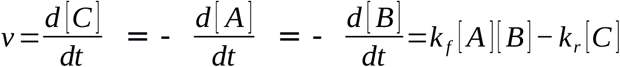

In order to reduce the number of rate constants that would need estimation, our aim was to use a minimal model with which we could still study the coincidence detection ability of the enzyme and capitalize on the predictions of the molecular simulations. We have used two versions of the model, one with the allosteric exclusion and the other with the simultaneous binding scheme for AC5 regulation in Fig. 3. The full reaction networks are given in Fig. S8. The two versions of the model have 8 and 16 rate constants, respectively. In Fig. 3, we have additionally used versions of the model which included cAMP production to illustrate the correspondence between k_c_ and the levels of cAMP and thus rationalize the use of k_c_ as a proxy for the cAMP levels. The reactions and parameters for cAMP production and degradation have been taken from [15].

There are no receptors included in the model, and the Da ↑ and ACh ↓ inputs are modeled as changes in the rate constants for the conversion of inactive to active G proteins. Pools of inactive G_olf_ and Gi are activated at rates of *k_fG_olf__*=5*s*^−1^ and *k_fG__i_*=5*s*^−1^ when Da or ACh is present. This eliminates the parameters that would have corresponded to the reactions of ligand and receptor binding, and G protein and receptor binding. Additionally, we have omitted the heterotrimeric nature of the G proteins, i.e. we have not included the Gβγ subunit in the model. The G proteins are modeled as simply switching between an active and inactive form. The value for the rate of Gα_i_ activation is chosen so that there is a high resting-level inhibition of AC5 by the tonic level of ACh, which is a plausible assumption based on recent experimental results [14]. The value for k_fG_olf__ is, similarly, chosen to achieve amounts of active G_olf_ high enough to drive the binding reactions with AC5 and AC5 · Gα_i_ forward. The total amounts of AC5 and the two G proteins used for this model are n_AC5_ = 1500 nM, n_Golf_ = 1500 nM and n_Gi_ = 6000 nM.

The activated Gα_olf_ and Gα_i_ interact with AC5 and can form each of the binary complexes and the ternary complex. Their deactivation is done by the intrinsic GTPase activity of the G proteins themselves, but is increased by AC5 for the case of Gα_s_ (a homologue to Gα_olf_) at least fivefold and the regulator of G protein signaling 9-2 (RGS9-2) for Gα_i_ 20 to 40 times [41, 79], for which reason we have used values of *k_rG_olf__*=0.5*s*^−1^ and *k_rG_i__*=5*s*^−1^. If deactivated, the G proteins unbind from AC5. For the reverse (unbinding) rates of the G proteins from the binary AC5 complexes, we use values 100 times greater than the corresponding forward rate constants, which is the order of magnitude fitted in [23]. The reverse rates of G protein unbinding from the ternary complex, are increased by an order of magnitude compared to the reverse rates for unbinding from the respective binary complexes to qualitatively incorporate possible reduced stability of the ternary complex compared to the binary complexes on longer time scales (see Results section).

The remaining parameters, the forward rate constants of the G proteins’ binding to AC5 and the binary complexes AC5 · Gα_olf_ and AC5 · Gα_i_, were estimated with the BD simulations (see Table 1). We have also varied these to explore their effects on the network’s ability to perform coincidence detection.

The kinetic model and related scripts to produce some of the figures can be found at https://github.com/danieltrpevski/AC5-kinetic-model.

### Measures of Coincidence Detection

As was defined in the introduction, for the signal transduction network that we consider, coincidence detection means to respond with significant amounts of cAMP only when the two incoming signals Da ↑ and ACh ↓ coincide in time and in the spatial vicinity of the receptors. Note that there are two aspects of coincidence detection in the definition:

a. distinguishing between the inputs Da ↑ + ACh ↓, and a Da ↑ or ACh ↓ alone, and
b. responding strongly enough with amounts of cAMP that are physiologically relevant.

To quantify how well the network distinguishes between the different inputs, we use the synergy quantity, and to quantify the strength of the response, we use the average catalytic rate, both defined below.

#### Average Catalytic Rate

The average catalytic rate is an average of the catalytic rates of the unbound form of AC5 and each of the complexes with the Gα subunits. For the allosteric exclusion scheme in Fig. 3B, where the ternary complex does not form, it is:

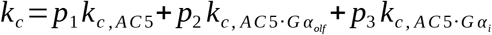

with the weights

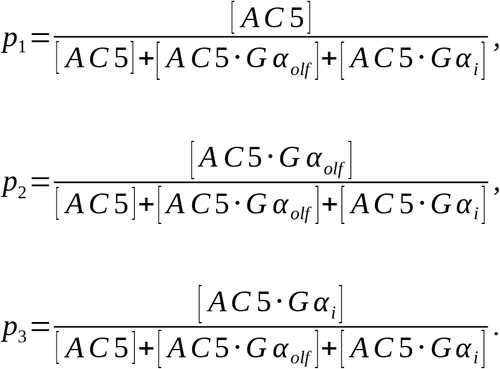

being the amounts of each enzyme species in the allosteric exclusion scheme as a fraction of the total concentration of AC5 in the system. For the simultaneous binding scheme where the ternary complex does form, the average catalytic rate is

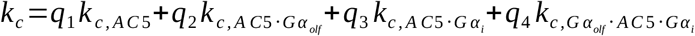

with

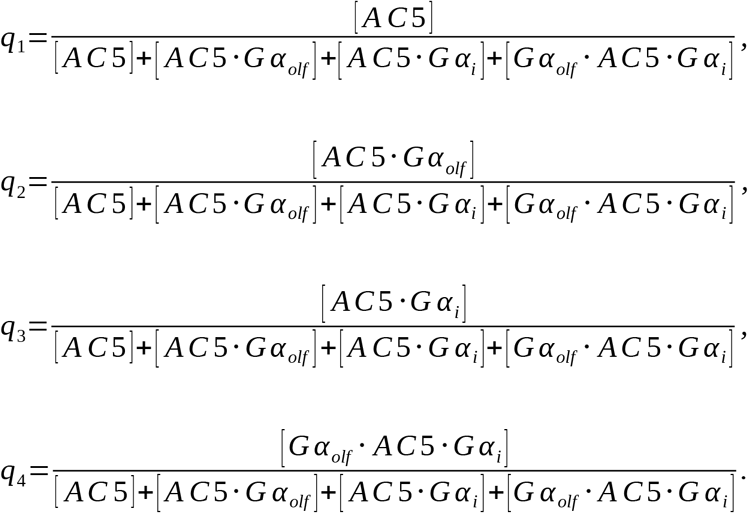

We assume that the catalytic rate of the unbound AC5 is *k*_*c,AC*5_=0.1*s*^−1^, and this rate is scaled by factors of *α_Gα_olf__* and *α_Gα_i__* when the respective regulator G protein subunit binds:

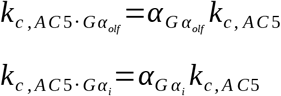

The factors of stimulation and inhibition of AC5 are set to be *α_Gα_olf__* = 200 ([23]) and *α_Gα_i__* =0.01*s*^−1^. For the catalytic rate of the ternary complex, we use the result of the MD simulations that the ternary complex is inactive, *α_Gα_olf_,Gα_i__*=*α_Gα_i__*, i.e.:

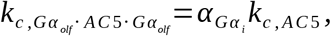

except in Figs. 3D, 3E and 3F, where the catalytic rate of the ternary complex is varied to investigate its effect on coincidence detection.

#### Synergy

The synergy of two signals s_1_ and s_2_ is defined as

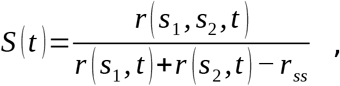

where r_ss_ is the response at steady state. This quantity measures how strong the response r of the signal transduction network is for two coincident signals compared to the responses for single signals. The synergistic effect of the input signals can be examined in light of any quantity of interest in the network that is affected by the inputs, such as the level of activated PKA, for example [15]. Not having included PKA or cAMP in the kinetic model, we use the average catalytic rate k_c_ as a proxy for the amount of cAMP produced, since the latter directly depends on k_c_. That is, the synergy of a simultaneous Da ↑ and ACh ↓ is:

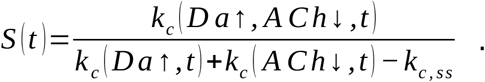

S(t) > 1 indicates a nonlinearly greater response in the presence of the two coincident signals, S(t) = 1 indicates a linear response to the coincident signals, and S(t) < 1 is a sublinear response. Hence, the case S(t) > 1 indicates that the signal transduction network can perform coincidence detection, whereas S(t) ≤ 1 indicates an inability to do so.

Example traces for k_c_ and the corresponding synergy are given in Fig. 6. Using

**Figure 6.**
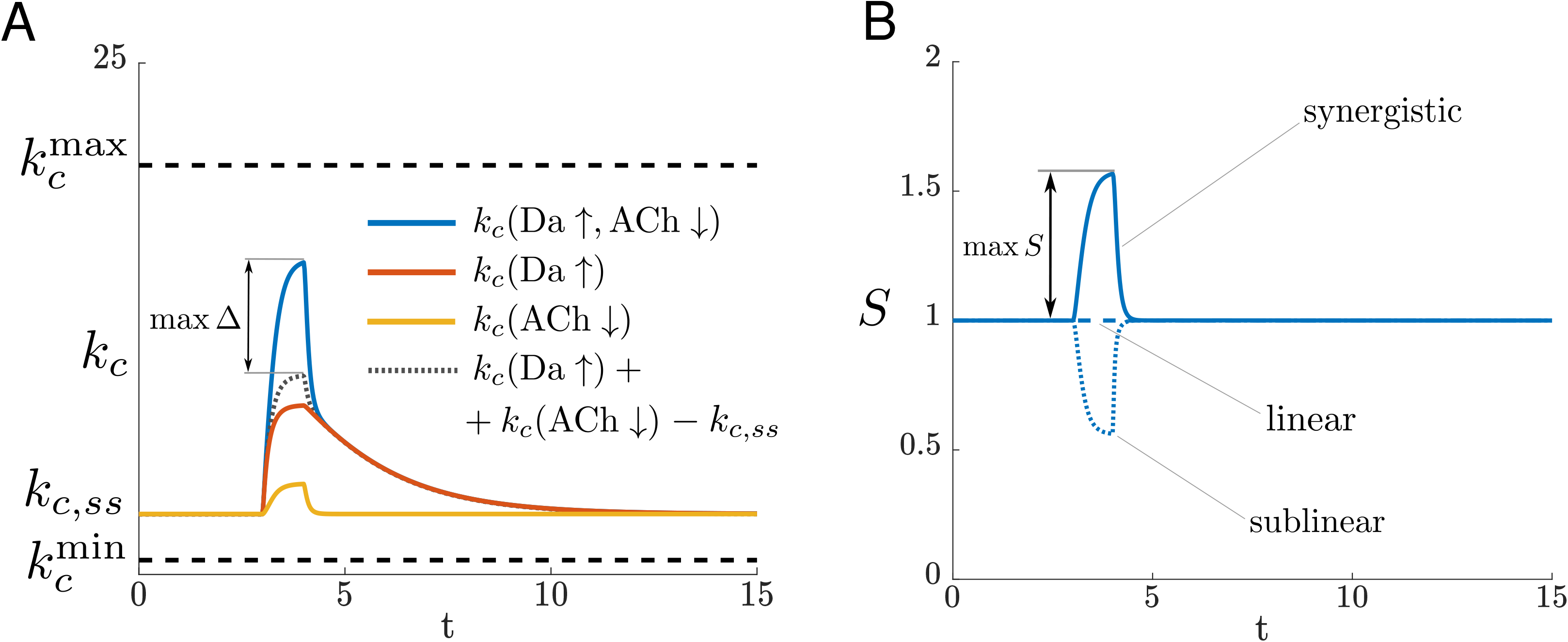
**A** A cartoon of the average catalytic rate during each of the input situations Da ↑ + ACh ↓, Da ↑, and ACh ↓. **B** The synergy corresponding to panel (A). In addition, the dashed and dotted lines illustrate how the synergy for a linear and a sublinear response would look like, respectively.

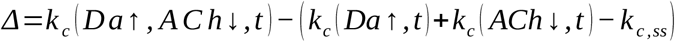

to express the difference between the response of the network for two coincident signals and the response for single signals, the expression for the synergy can also be rewritten as

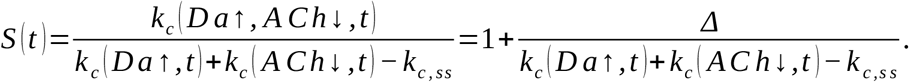

The difference in the responses, Δ, determines how big the synergistic effect of the input signals is (Fig. 6).

Figure 6 is an example depicting how k_c_ relates to the synergy. There are minimum and maximum bounds on k_c_: it would attain the minimum value k_c_^min^ if all of AC5 were inhibited by Gα_i_, that is, only the catalytically inactive complex AC5 · Gα_i_ exists, where k_c_^min^ is the catalytic rate of AC5 · Gα_i_ (see above). Analogously, it would attain the maximum value k_c_^max^ if all of AC5 were bound in the catalytically active complex AC5 · Gα_olf_, where k_c_^max^ is its catalytic rate (see above). In the models we use in this study, AC5 is never fully occupied by either of the Gα subunits, and hence k_c_ is always between the minimum and maximum bounds. To maximize *Δ*, one would need to have k_c_ grow towards the maximum as much as possible during Da ↑ + ACh ↓, and to have it as low as possible at steady state, and when each of the signals Da ↑ and ACh ↓ comes alone.

#### The metric C combines S and k_c_

A combination of the average catalytic rate and the synergy gives a metric which can be helpful to evaluate whether the network both distinguishes between the different input situations and responds strongly. We use a product of k_c_ and S for this purpose, *C*=*Sk_c_*.

## Acknowledgements

This research has received funding from the European Union Seventh Framework Programme (FP7/2007-2013) under grant agreement N° 604102 (HBP) (JHK, PC, RCW), the European Union’s Horizon 2020 Framework Programme for Research and Innovation under the Specific Grant Agreement N°s 720270 (Human Brain Project SGA1) (JHK, PC, RCW, UR, AGN) and 785907 (Human Brain Project SGA2) (JHK, PC, RCW, UR, DT), the Swedish e-Science Research Centre (SeRC) (JHK, DT), the Swedish Research Council (JHK, AGN, DT), the Klaus Tschira Stiftung (RCW, NJB), National Institute on Alcohol Abuse and Alcoholism Grant 2R01AA016022 (JHK). The authors thank Pietro Vidossich for scientific discussions during this project.

## Author contributions

PC, UR, RCW, and JHK conceived the study. AGN, NJB, DN, DT, SCvK, PC, RCW, UR and JHK designed the study. DN set up and performed the molecular dynamics simulations of ternary complexes and the *apo* AC5 · Gαolf complex. SCvK prepared the starting structures of all MD simulations, set up and performed the MD simulation of the *apo* AC5 · Gα_i_ complex and helped setting up the simulations of the ternary complexes. DN and SCvK analyzed MD simulations. NJB performed and analyzed the Brownian dynamics simulations. DT set up and performed the kinetic modeling and analyzed the results. SCvK and NJB performed the isoform sequence analysis. All authors analyzed the data. DT, DN, NJB and SCvK wrote the manuscript. AGN, RCW, PC, UR and JHK reviewed and revised the manuscript.

## Supplementary Information

**Figure S1:**
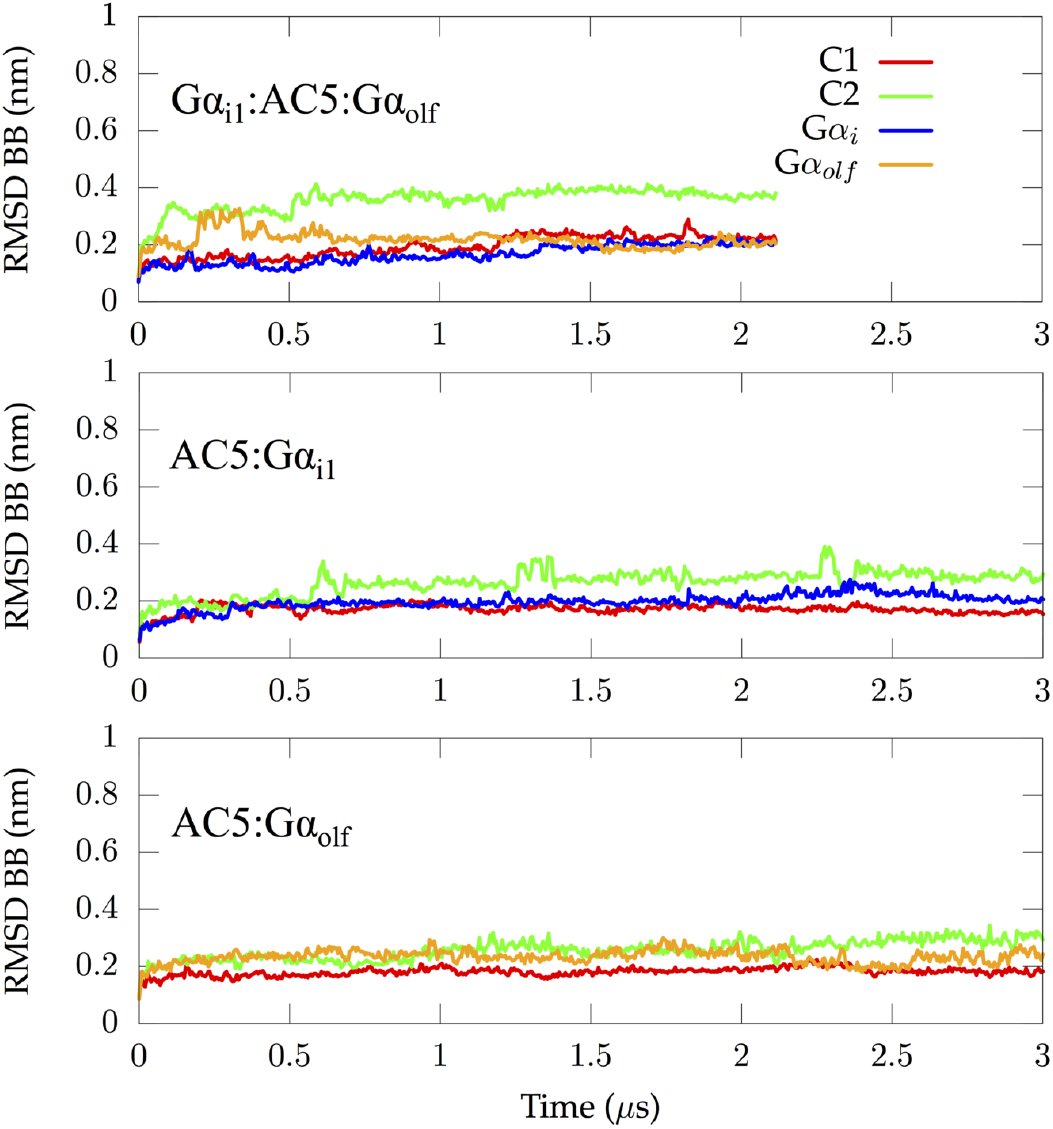
Root-mean-square deviations of three *apo* complexes: Gα_olf_ · AC5 · Gα_i1_, AC5 · Gα_i1_ and AC5 · Gα_olf_. (Top panel) Individual RMSD values for C1, C2, Gα_i1_ and Gα_olf_ in the ternary complex only including the backbone of the domains. In the case of Gα_i1_, residues 33 to 345 were taken into account during the RMSD calculation as the C and N termini are not structured. (Middle panel) Individual RMSD values for C1, C2 and Gα_i1_ in the binary AC5 · Gα_i1_ complex only including the backbone of the domains. In the case of Gα_i1_, residues 33 to 345 were taken into account during the RMSD calculation as the C and N termini are not structured. (Bottom panel) Individual RMSD values for C1, C2 and Gα_olf_ in the binary AC5 · Gα_olf_ complex only including the backbone of the domains.

**Figure S2:**
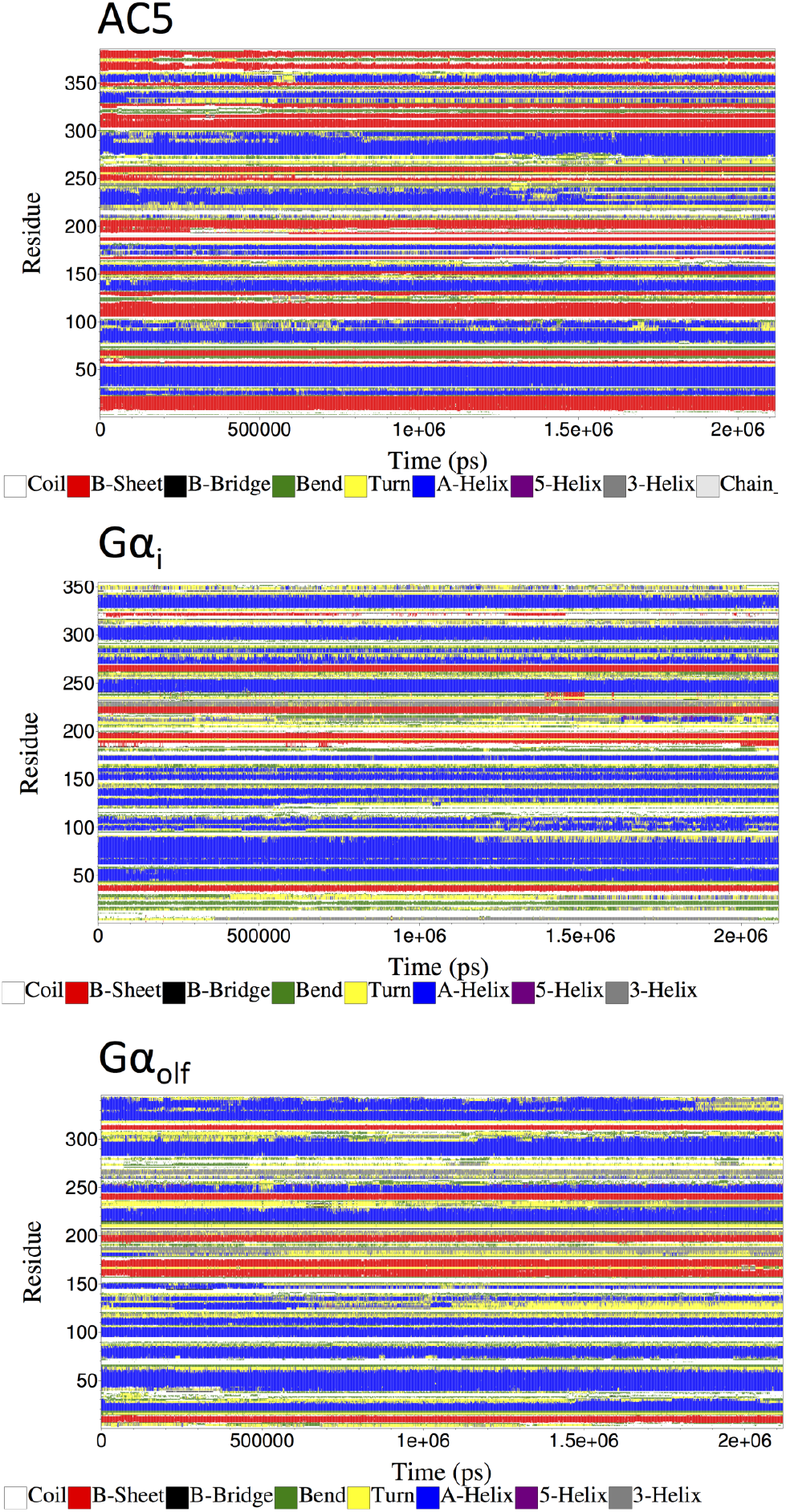
Time evolution of the secondary structures for AC5 (top), Gα_i_ (middle), and Gα_olf_ (bottom) along the trajectory of the *apo* ternary complex.

**Figure S3:**
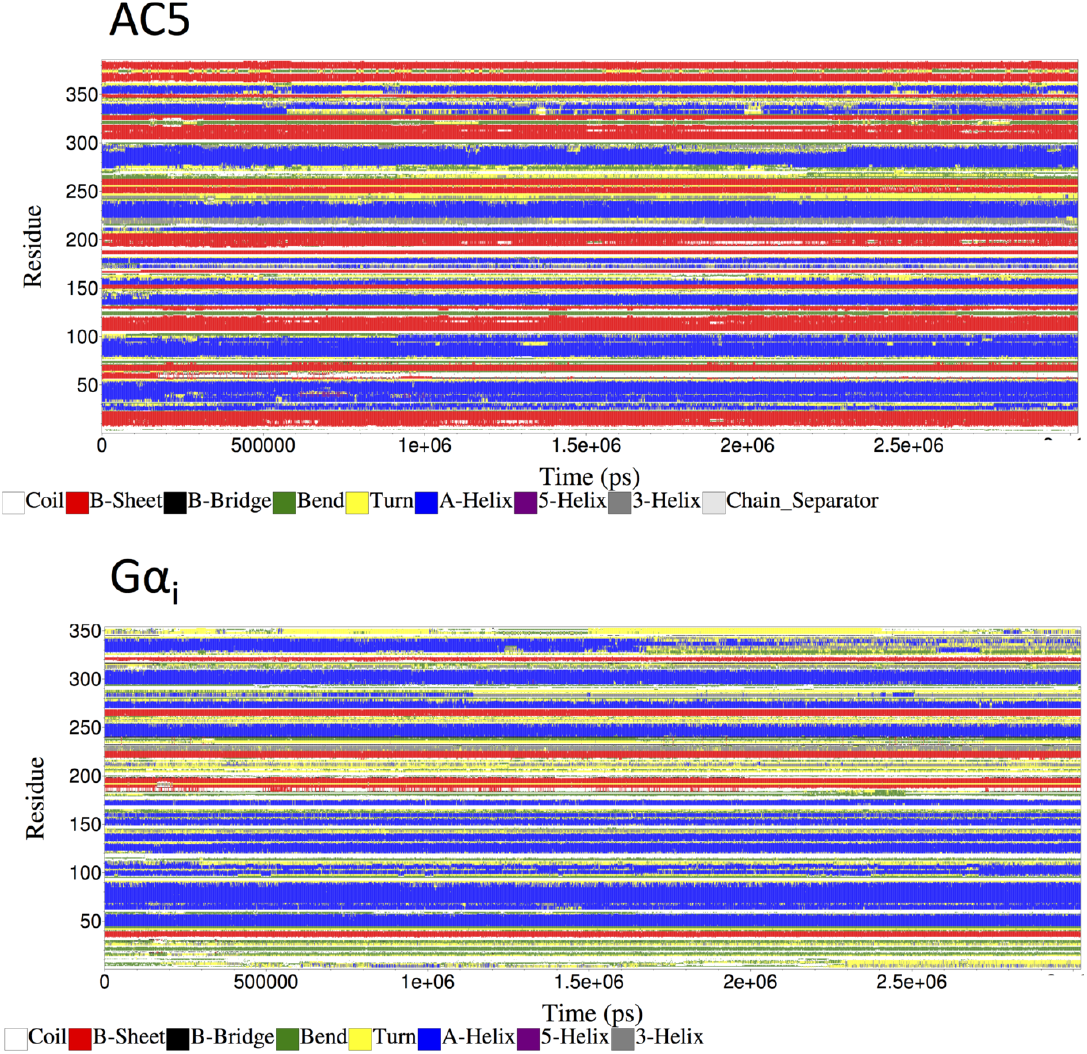
Time evolution of the secondary structures for AC5 (top), and Gα_i_ (bottom), along the trajectory of the *apo* binary complex AC5 · Gα_i_.

**Figure S4:**
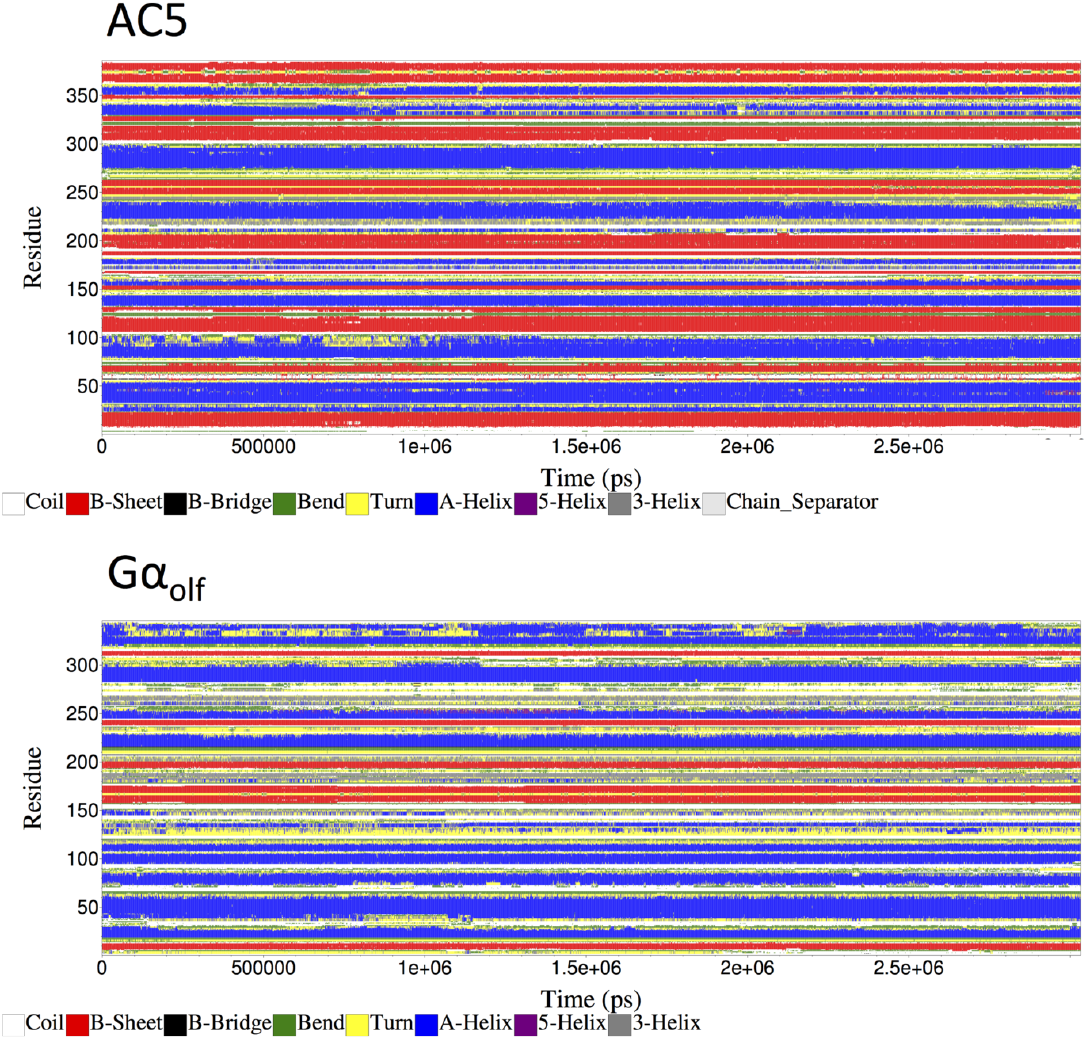
Time evolution of the secondary structures for AC5 (top), and Gα_olf_ (bottom), along the trajectory of the *apo* binary complex AC5 · Gα_olf_.

**Figure S5:**
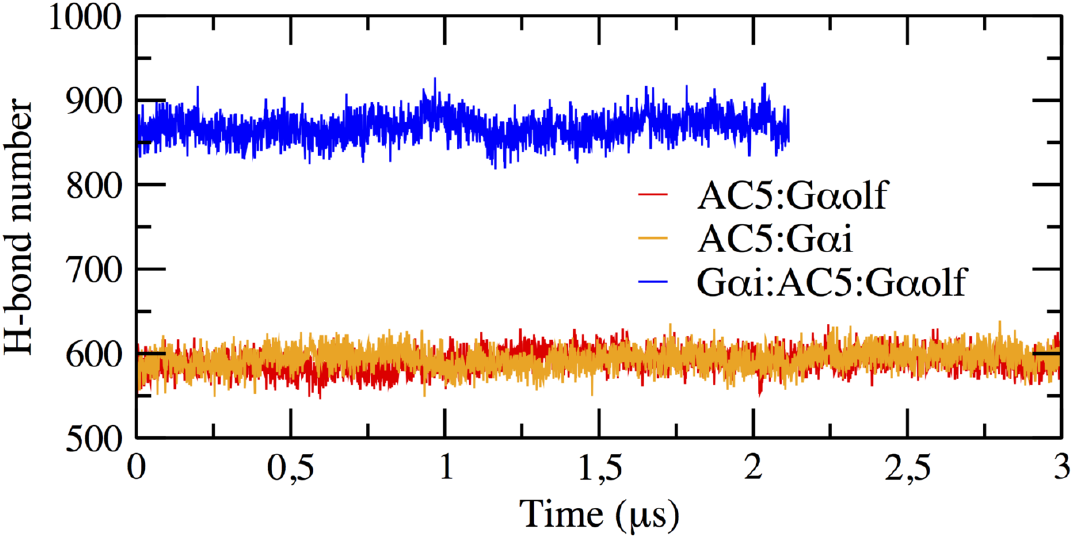
Time evolution of the number of hydrogen bonds present in the three simulated *apo* complexes along the respective MD trajectories.

**Figure S6:**
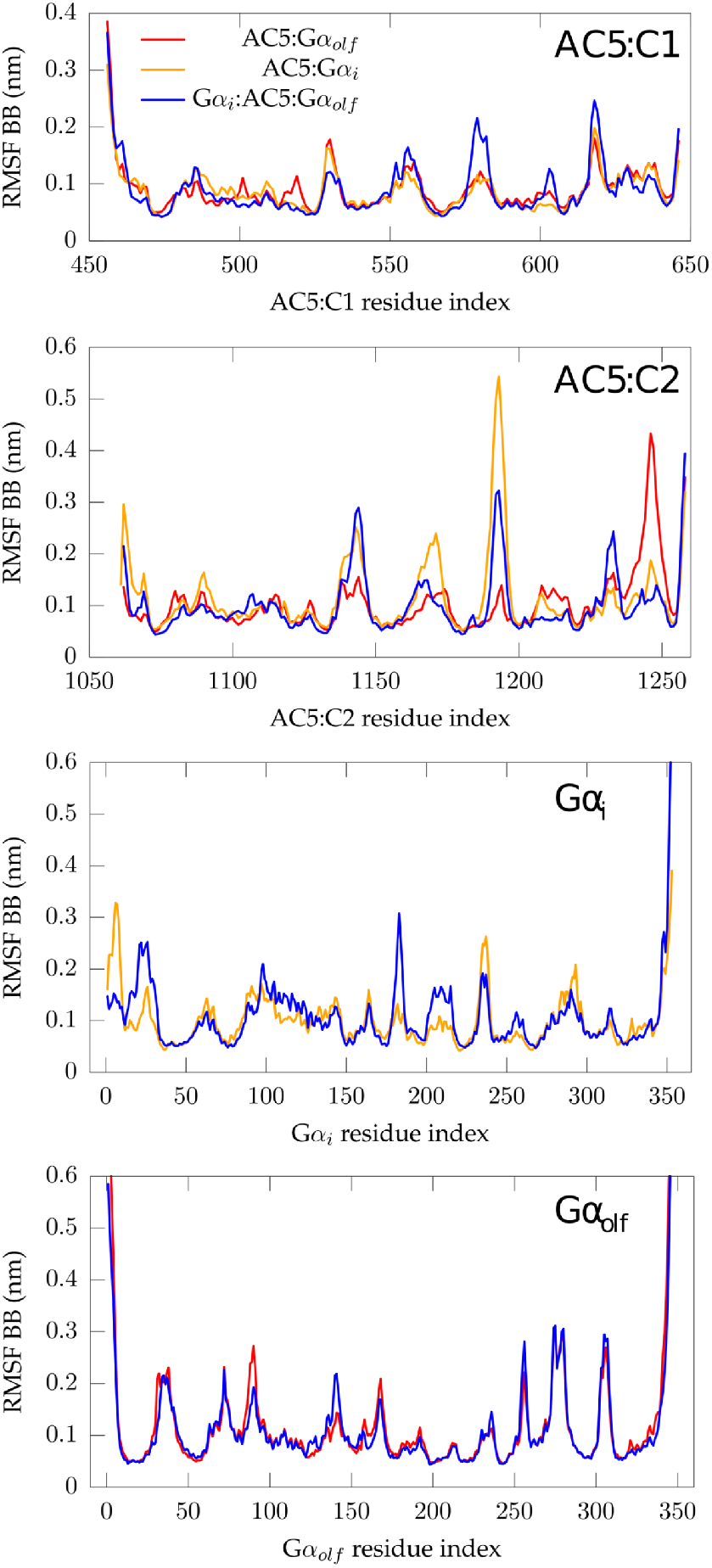
Root-mean-square fluctuations per residue calculated on the protein backbone of the different subunits (from top to bottom, AC5:C1, AC5:C2, Gα_i_, and Gα_olf_) of the three simulated *apo* complexes.

**Figure S7:**
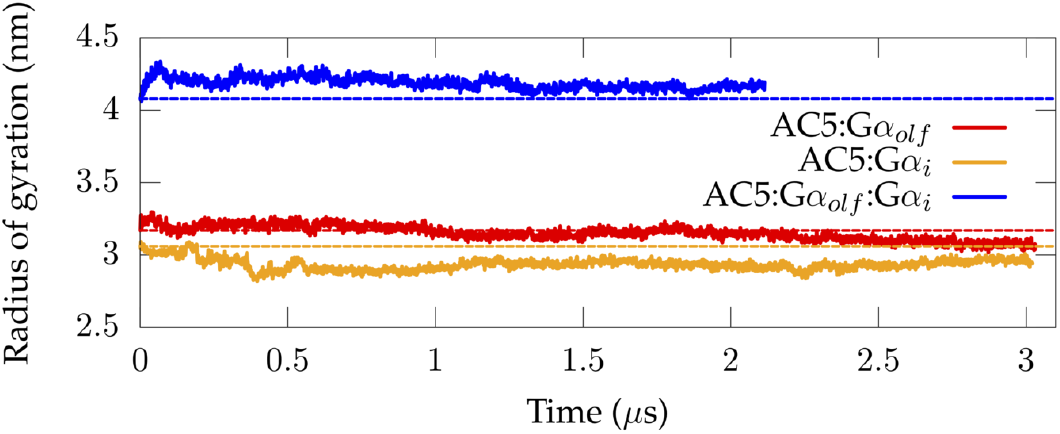
Radius of gyration calculated along the MD trajectories of the three simulated *apo* complexes, Gα_olf_ · AC5 · Gα_i_, AC5 · Gα_i_, and AC5 · Gα_olf_. The dashed lines indicate the values of the radius of gyration in the initial

**Figure S8:**
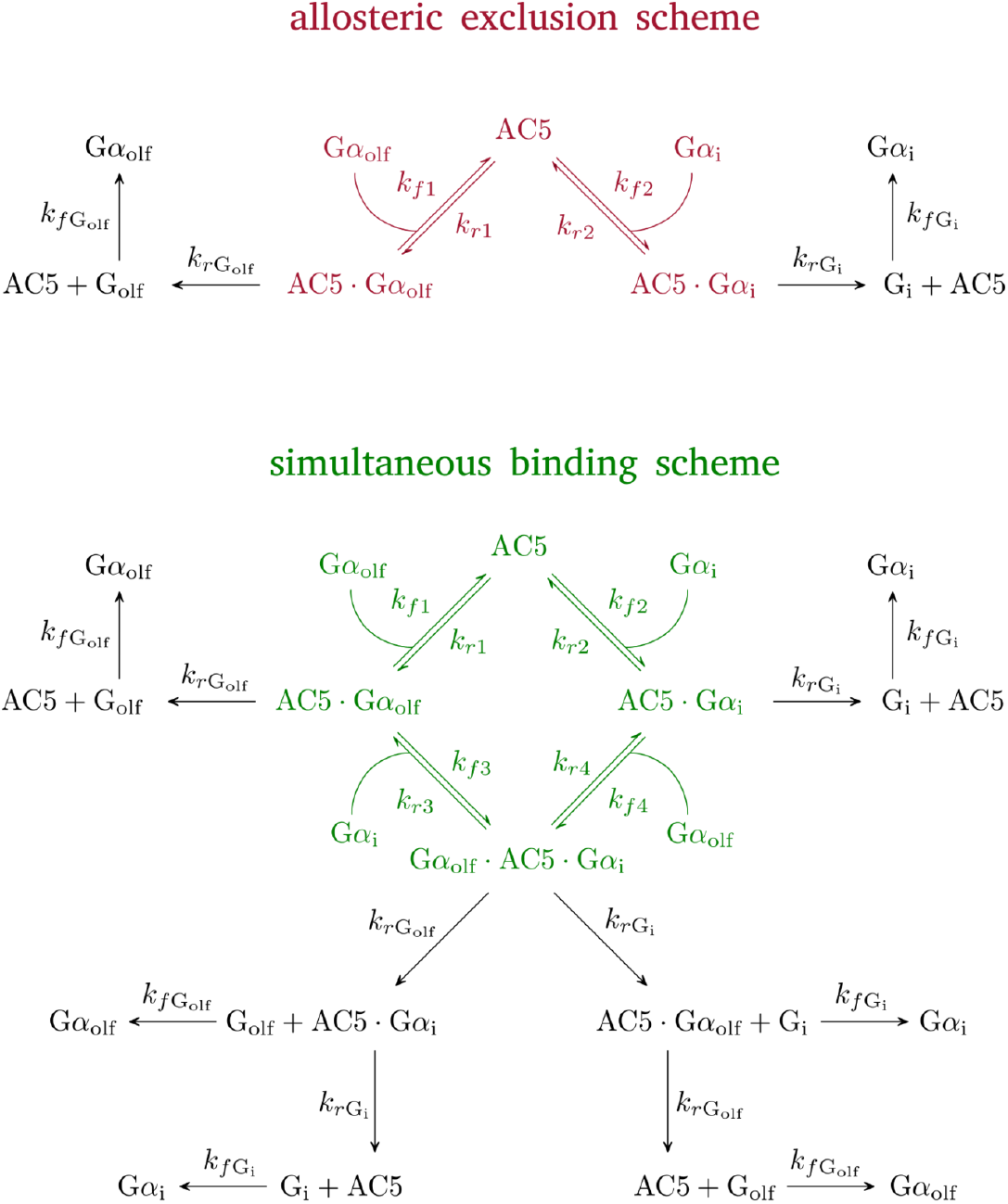
The full kinetic models of the signal transduction networks used in this study.

**Figure S9:**
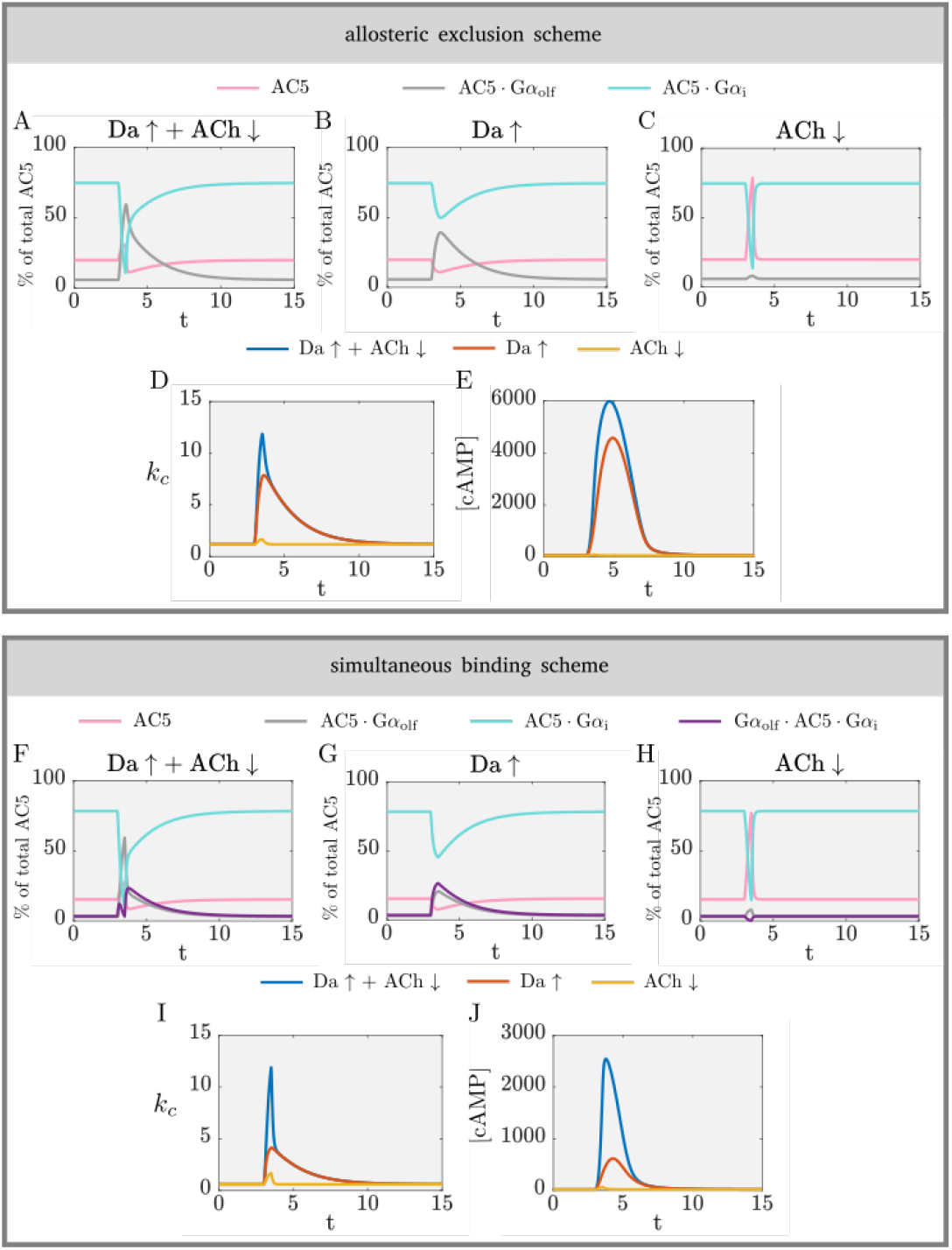
The effect of the interaction motif between AC5 and the regulatory Gα subunits on coincidence detection. For the allosteric exclusion and simultaneous binding schemes, respectively, the amounts of each enzyme species as a percentage of the total amount of AC5 are shown for the cases of Da ↑ + ACh ↓ (A, F), Da ↑ (B, G), and ACh ↓ (C, H). (D, I) Average catalytic rate for each scheme. (E, J) cAMP levels for each scheme.

**Figure S10:**
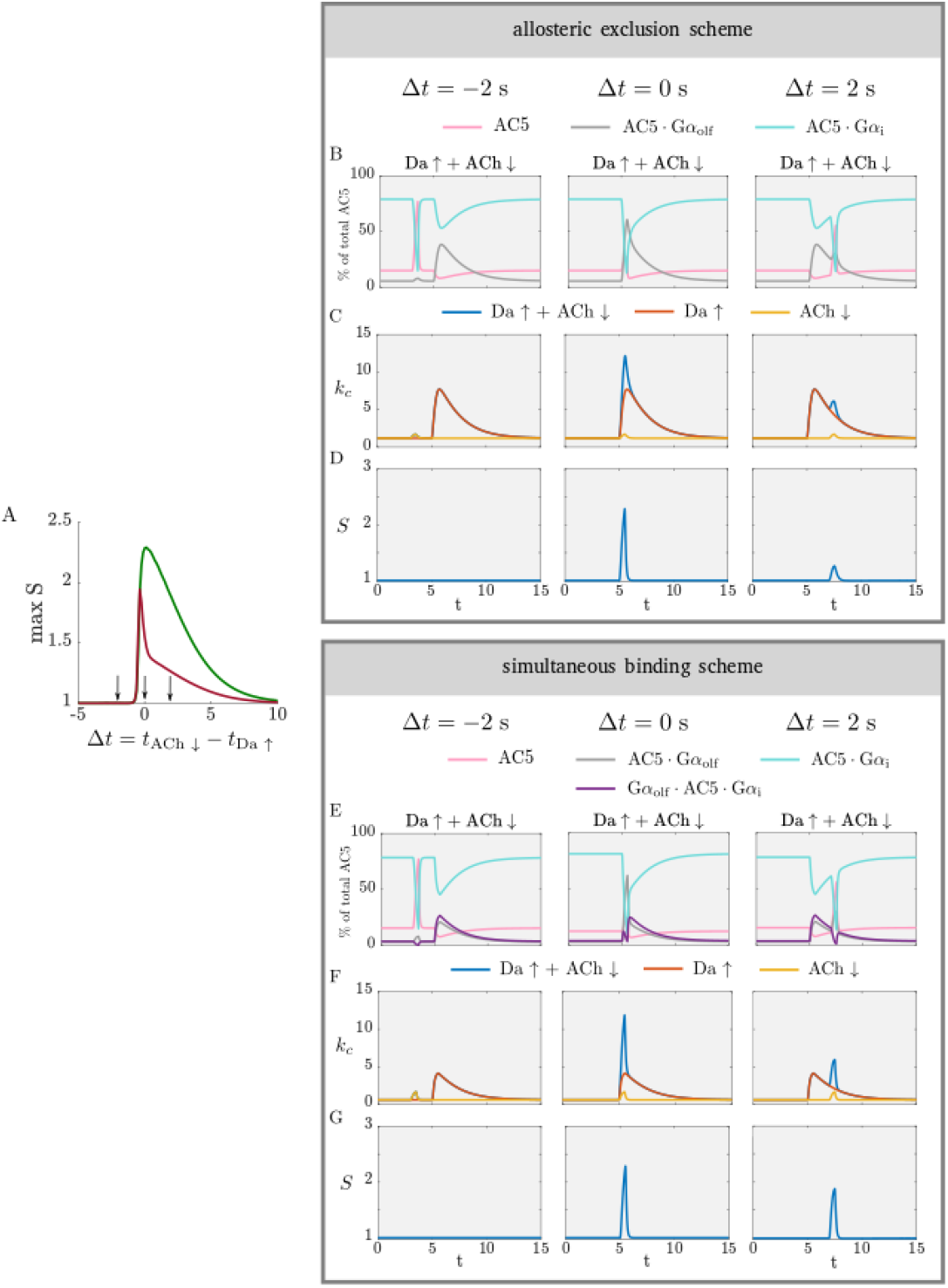
(A) The detection window for the allosteric exclusion scheme and simultaneous binding scheme. Arrows are the time differences between ACh ↓ and Da ↑ chosen for the traces below. (B) The percentage of each AC5 species as a fraction of the total amount of AC5, (C) average catalytic rate, (D) synergy for the allosteric exclusion scheme. (E), (F), and (G) are the same quantities for the simultaneous binding scheme. Note the shared axes.

**Figure S11:**
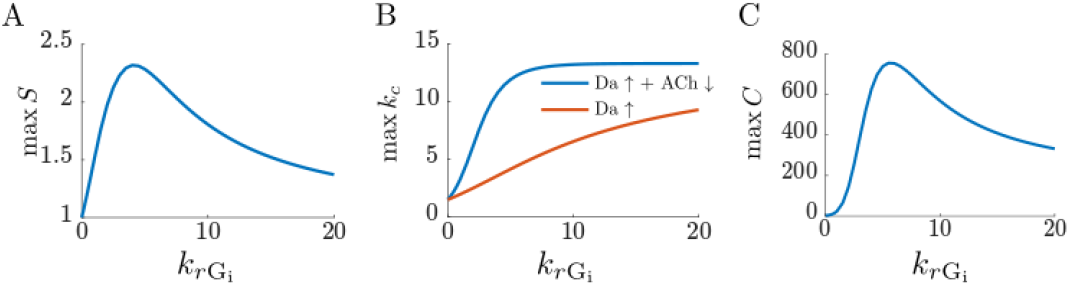
(A) The maximum of the synergy, (B) the maximum of k_c_, (C) the maximum of the metric C as dependent on the rate of Gα_i_ deactivation, k_rGi_.

**Figure S12:**
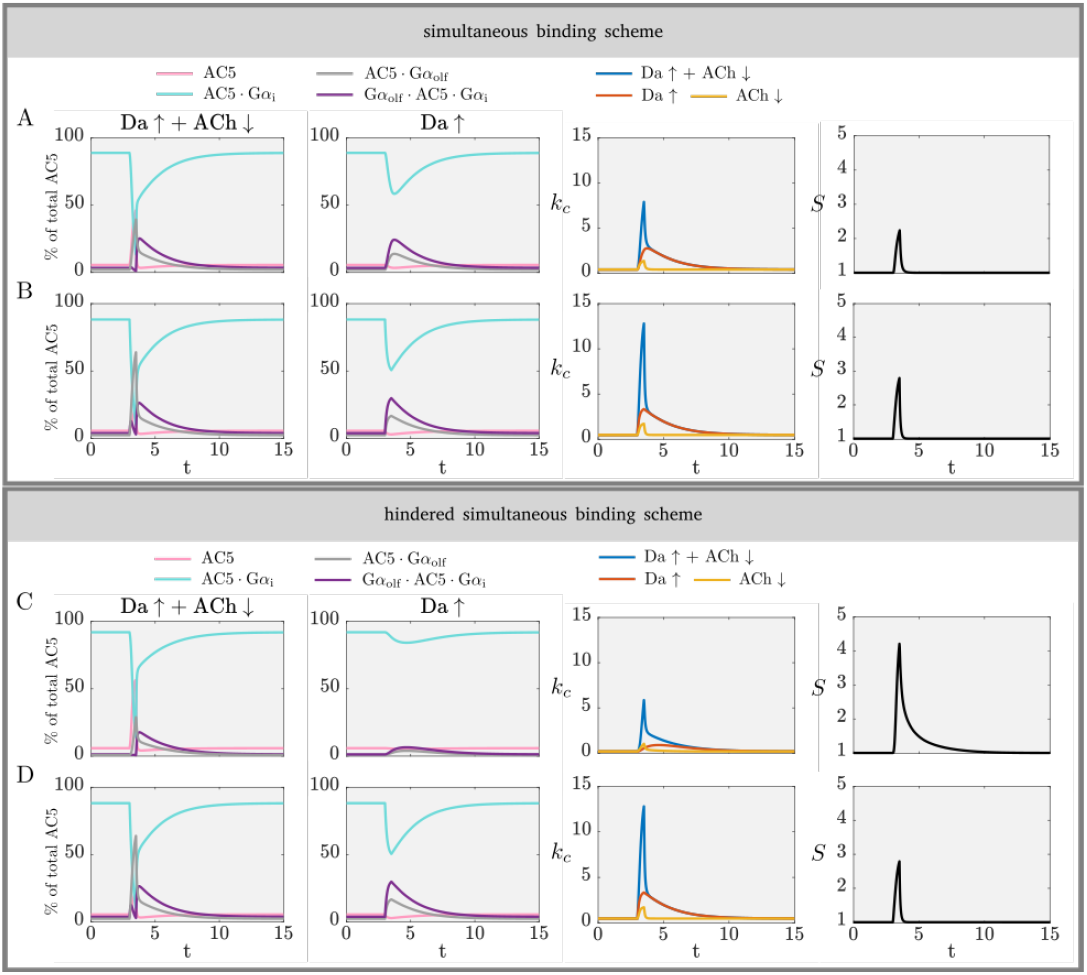
From left to right: the percentage of enzyme species for Da ↑ + ACh ↓, the percentage of enzyme species for Da ↑, the average catalytic rate, and the synergy, for the simultaneous binding scheme for (A) k_f1_ = k_f4_ = 0.002 (nMs)^−1^, k_f2_ = k_f3_ = 2 (nMs)^−1^, and (B) k_f1_ = k_f4_ = 2 (nMs)^−1^, k_f2_ = k_f3_ = 2 (nMs)^−1^ and the hindered simultaneous binding scheme for (C) k_f1_ = k_f4_ = 0.002 (nMs)^−1^, k_f2_ = k_f3_ = 2 (nMs)^−1^ and (D) k_f1_ = k_f4_ = 2 (nMs)^−1^, k_f2_ = k_f3_ = 2 (nMs)^−1^. Note the shared axes.

**Figure S13:**
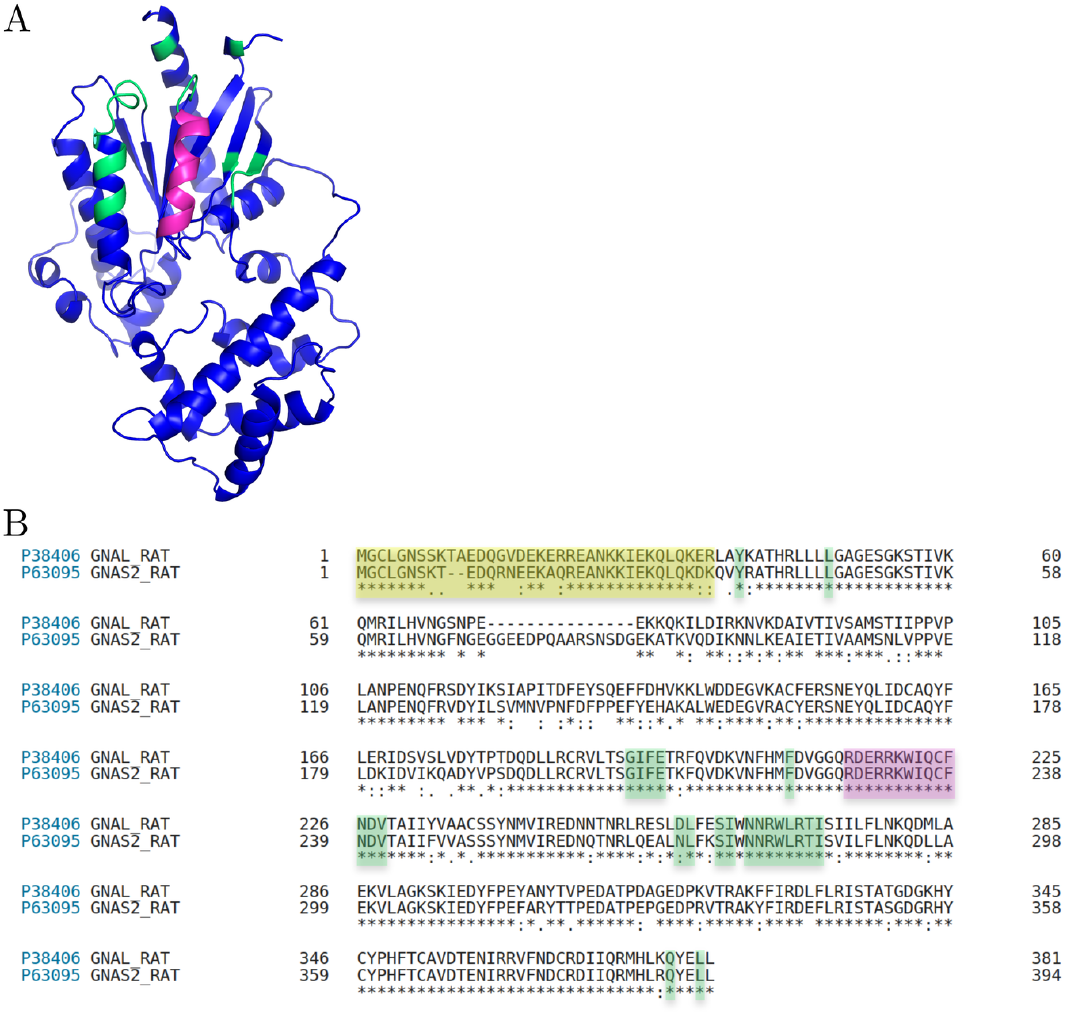
The structure of Gα_olf_ in the modelled AC5 · Gα_olf_ *apo* complex (A). The highlighted regions show the switch II helix residues that interact the C2 binding groove on AC5 (magenta) and other amino acid residues that are within 6 of AC5 in the modelled structure. The sequence alignment of rat Gα_olf_ (GNAL) and Gα_s_ (GNAS2) (B). The magenta and green regions show the residues highlighted in (A). The yellow region shows the N-terminal residues not included in the structure used in this work.

**Figure S14:**
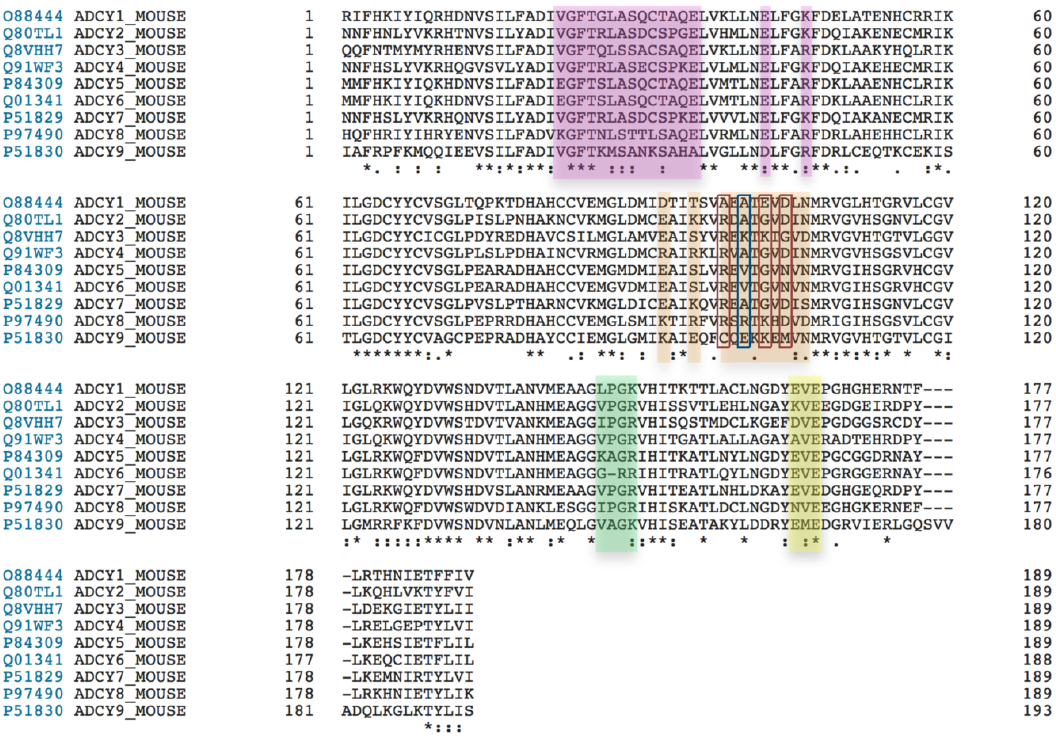
Multiple sequence alignment for all mouse AC isoforms with the colors matching those the structure in Fig. 5A. The sequences were taken from Uniprot, and aligned using Clustal Omega within Uniprot. The red and blue boxes show positions where AC1 has substitutions compared to AC5, as described in Fig. 5.

**Figure S15:**
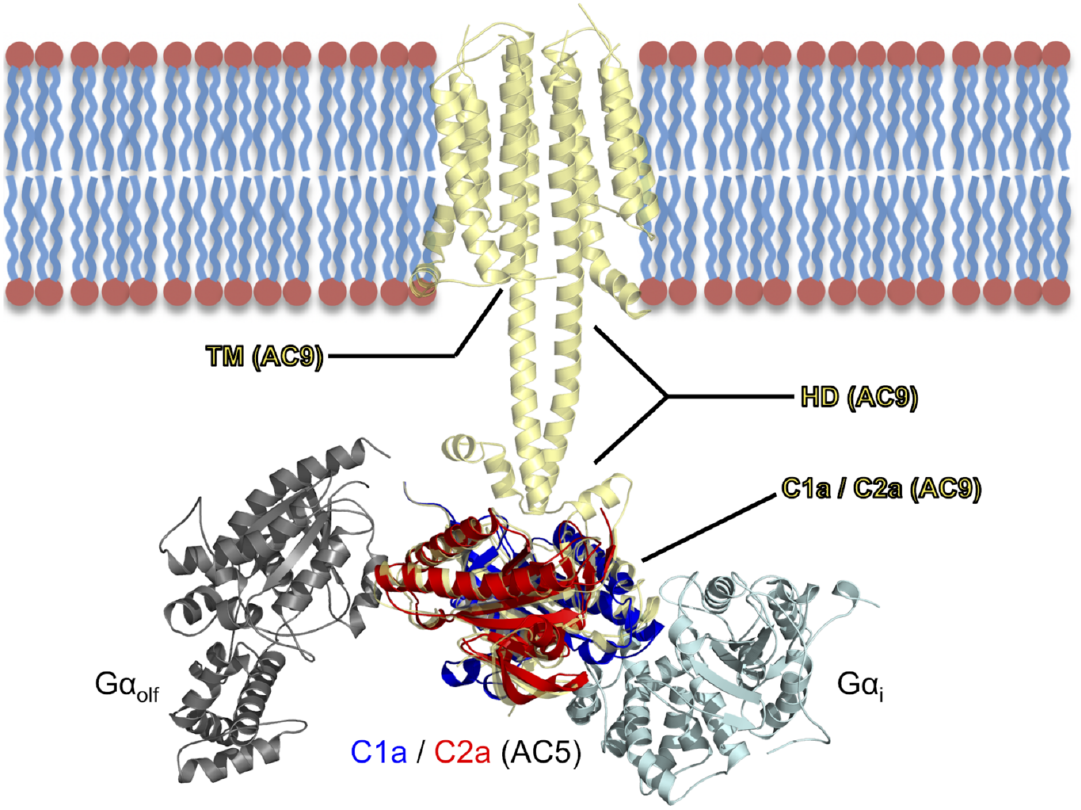
Initial modelled configuration of the Gα_olf_ · AC5 · Gα_i_ ternary complex used in our classical MD simulations, after fitting of Cα atoms of C1a and C2a domains on the respective Cα atoms of the AC9 protein (PDB ID: 6R3Q). AC9 (pale yellow) consisting of the transmembrane domain (TM), the helical domain (HD), and the two pseudo-symmetric C1a and C2a domains, is fully shown in cartoon representation. The two domains C1a (blue) and C2a (red) of AC5 in complex with Gα_i_ (cyan) and Gα_olf_ (gray) proteins are also represented as a cartoon.

**Table S1.** Bimolecular association rate constants (nMs)^−1^ for the forward reactions computed via BD simulations. Each rate constant was calculated using a number of snapshots from MD simulations. The reported numbers for each snapshot are the mean values estimated from 4 BD simulations of 50 000 trajectories (standard deviation in parentheses). Rate constants were calculated for complexes including *apo* and *holo* AC5 (superscripts a and h respectively).

| MD snapshot | $k_{f1}^a$ | $k_{f1}^h$ | $k_{f2}^a$ | $k_{f2}^h$ |
| --- | --- | --- | --- | --- |
| 1 | 0.003 (0.002) | 0.013 (0.004) | 0.015 (0.002) | 0.005 (0.002) |
| 2 | 0.011 (0.003) | 0.045 (0.007) | 0.018 (0.006) | 0.028 (0.004) |
| 3 | 0.038 (0.003) | 0.019 (0.002) | 0.036 (0.006) | 0.10 (0.01) |
| 4 | 0.022 (0.006) |  | 0.008 (0.003) | 0.061 (0.006) |
| 5 |  |  | 0.016 (0.003) |  |
| 6 |  |  | 0.013 (0.003) |  |
| 7 |  |  | 0.027 (0.005) |  |
| Mean | 0.018 (0.007) | 0.026 (0.009) | 0.02 (0.01) | 0.05 (0.01) |

| MD snapshot | $k_{f3}^a$ | $k_{f3}^h$ | $k_{f4}^a$ | $k_{f4}^h$ |
| --- | --- | --- | --- | --- |
| 1 | 0.009 (0.003) | 0.004 (0.001) | 0.037 (0.004) | 0.035 (0.004) |
| 2 | 0.004 (0.001) | 0.017 (0.004) | 0.007 (0.003) | 0.016 (0.003) |
| 3 | 0.011 (0.005) | 0.032 (0.004) | 0.007 (0.003) | 0.013 (0.002) |
| 4 | 0.005 (0.002) | 0.016 (0.002) | 0.0087 (0.0007) | 0.016 (0.003) |
| 5 | 0.030 (0.004) |  | 0.06 (0.01) |  |
| 6 |  |  |  |  |
| 7 |  |  |  |  |
| Mean | 0.012 (0.008) | 0.017 (0.006) | 0.02 (0.01) | 0.020 (0.006) |

### The allosteric exclusion scheme inherently lacks the ability for coincidence detection

To show this we make the following simplifications:

1. During a Da ↑ and a ACh ↓, respectively, the rise in [Gα_olf_] and drop in [Gα_i_] are assumed to be square-shaped, i.e. the interactions between the G proteins and AC5 are assumed to be sufficiently fast so that the transient steady state levels in the network are achieved quickly and last for the whole duration of the signals
2. During the Da ↑, enough Gα_olf_ is produced to occupy all AC5, i.e. [Gα_olf_] ≫ [Gα_i_] and [Gα_olf_] is at a saturating level.
3. During the ACh ↓, [Gα_i_] drops to approximately 0, i.e. all available AC5 is disinhibited from Gα_i_;

We consider what happens in the limiting cases of ‘perfect’ stimulation and inhibition of AC5, i.e. *α*_G_olf__ → ∞ and *α*_G_i__ = 0. With these assumptions only the effect of the structure of the regulatory scheme on coincidence detection is isolated.

We now determine the synergy of this scheme. The expression for the synergy is repeated here for convenience:

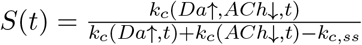

Each of the terms in the expression for the synergy are as follows. For Da ↑ + ACh ↓, there is no Gα_i_ in the system during a ACh ↓, so all AC5 is occupied by the produced saturating concentration of Gα_olf_.

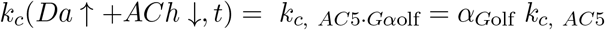

For Da ↑ alone, there is enough Gα_olf_ to outcompete Gα_i_ in the occupation of AC5, and AC5 is saturated with Gα_olf_, which yields the same result as for k_c_ (Da↑ + ACh↓):

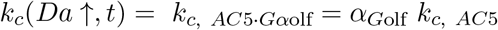

For ACh ↓ alone, there is no Gα_i_ in the system and AC5 is partly occupied by any resting-state levels of Gα_olf_:

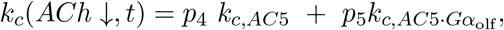

where

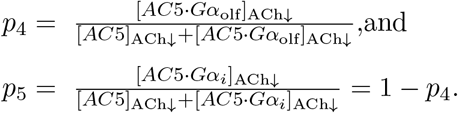

The steady state value for k_c_ is given under the section ‘Average catalytic rate’ in the Methods of the main text. Substituting these expressions in the expression for the synergy, and taking the limit *α*_G_olf__ → ∞ yields:

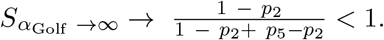

The synergy is always less than 1 since *p*_5_ > *p*_2_, i.e. *p*_5_ is the proportion of AC5 · Gα_olf_ when there is no Gα_i_ in the system, which is always greater than *p*_2_, the proportion of AC5 · Gα_olf_ in the resting state when there is G_i_ in the system.

An example of this result is given in Fig. S16, where the concentration of G_olf_ has been made very high to mimic the conditions used for the mathematical derivation. The duration of the signals is also very long to make the (transient) steady state levels evident. As shown in Fig. S16A and S16B, in the case of a Da ↑ the high amount of Gα_olf_ indeed outcompetes almost all of Gα_i_ in occupying the available AC5, and this input produces a similar average k_c_ as the case of Da ↑ + ACh ↓. The synergy in this scenario is less than 1 (Fig. S16F). However, the allosteric exclusion scheme can be used for coincidence detection with suitable choices in the amounts of the G proteins (given a set of rate constants for binding and unbinding). This is shown in Fig. 3B of the main text and its corresponding Fig. S9, where the amount of G_olf_ is not enough to occupy all available AC5.

**Figure S16:**
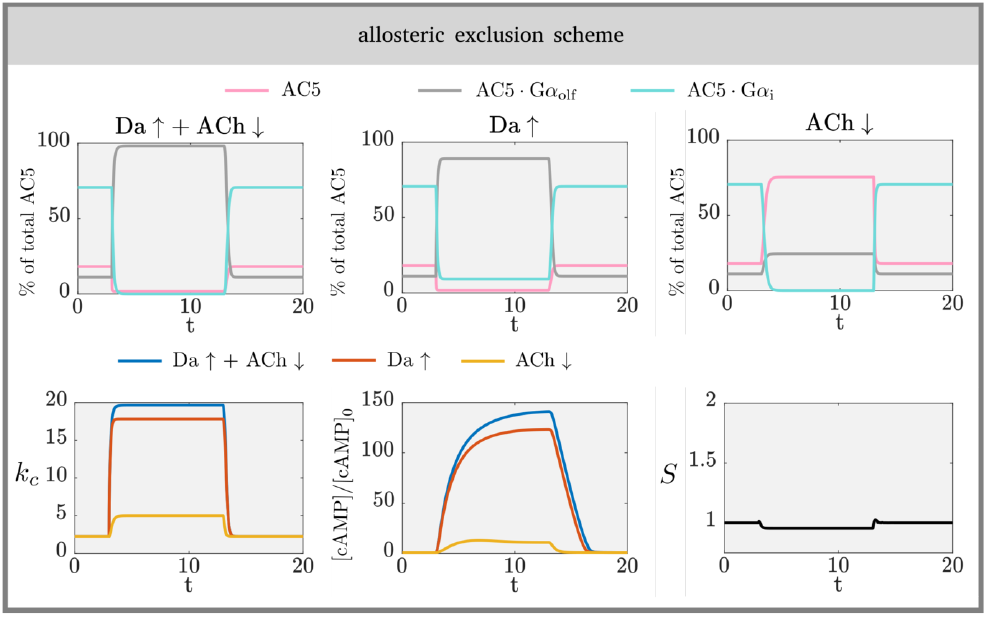
The allosteric exclusion scheme does not support coincidence detection for saturating concentrations of Gα_olf_.

